# Novel microRNAs targeting NMDA receptor subunits in animal models of schizophrenia

**DOI:** 10.1101/2021.10.04.463021

**Authors:** Sowmya Gunasekaran, Reena S. Jacob, Ramakrishnapillai V. Omkumar

## Abstract

N-methyl-D-aspartate receptors (NMDAR) are downregulated in schizophrenia possibly through microRNAs (miRNAs) that are differentially expressed in this condition. We screened the miRNAs that are altered in schizophrenia against the targets, *Grin2A* and *Grin2B* subunits of NMDAR using bioinformatic tools. Among the predicted miRNAs some interacted with the 3’-UTR sequences of *Grin2A* (miR-296, miR-148b, miR-129-2, miR-137) and *Grin2B* (miR-296, miR-148b, miR-129-2, miR-223) in dual luciferase assays. This was supported by downregulation of the GluN2B protein in primary hippocampal neurons upon overexpressing *Grin2B* targeting miRNAs. In two models of schizophrenia-pharmacological MK-801 model and neurodevelopmental methylazoxymethanol acetate (MAM) model which showed cognitive deficits - protein levels of GluN2A and GluN2B were downregulated but their transcript levels were upregulated. MiR-296-3p, miR-148b-5p and miR-137 levels showed upregulation in both models which could have interacted with *Grin2A*/*Grin2B* transcripts resulting in translational arrest. In MAM model, reciprocal changes in the expression of the 3p and 5p forms of miR-148b and miR-137 were observed. Expression of neuregulin 1 (*NRG1*), *BDNF* and *CaMKIIα*, genes implicated in schizophrenia, were also altered in these models. This is the first report of downregulation of GluN2A and GluN2B by miR-296, miR-148b and miR-129-2. Mining miRNAs regulating NMDA receptors might give insights into the pathophysiology of this disorder, providing avenues in therapeutics.

## 1. Introduction

Neurological disorders are among the most challenging contributors to global disease burden. Neuropsychiatric disorders majorly affect the adolescents and young adults globally [1]. Schizophrenia is a severe and complex neuropsychiatric disorder affecting 1% of the population worldwide [2]. It begins in the early age of 16-25 years and is considered as a neurocognitive disorder with neurodevelopmental origin [3]. It involves distortion of thinking and perception characterized by impaired cognition, positive psychotic symptoms including hallucinations, delusions, disorganized behavior and negative symptoms such as social withdrawal and apathy. Its etiology is unclear with suspected genetic and environmental factors. A major theory regarding the pathophysiology of schizophrenia is the glutamate hypothesis in general and NMDA receptor (NMDAR) hypothesis [4,5] in particular. This was proposed based on the observation that the psychotomimetic agents, phencyclidine (PCP) and ketamine, which are also NMDAR antagonists, induce negative and positive psychotic symptoms and neurocognitive disturbances similar to those found in schizophrenia by blocking neurotransmission through NMDARs [6–8]. PCP and ketamine block NMDARs in a noncompetitive fashion and other agents such as dizocilpine (MK-801), bind to a site located within the ion channel formed by the NMDAR complex. Clinical studies suggested that NMDAR hypofunction induced by PCP or ketamine may lead to secondary dopaminergic dysregulation in striatal and prefrontal brain regions [6]. Evidence from both animal and human studies suggest that the hyperdopaminergia associated with schizophrenia may, in fact, result from underlying dysfunction of NMDAR-mediated feedback mechanisms [9]. There has also been reports where dysfunction in calcium/calmodulin (CaM)-dependent protein kinase II (CaMKII) expression leads to dysregulation in the glutamatergic pathways by affecting NMDARs [10]. Although the exact mechanism on how abnormal CaMKII has a role in schizophrenia is not elucidated, it is noted that mutant animals lacking densin-180, the CaMKII-anchoring protein, have behavioural phenotypes similar to schizophrenia [10].

There are other candidate genes which are affected in schizophrenia such as Neuregulin 1 (NRG1) and brain-derived neurotrophic factor (BDNF) [11]. NRG1 is a developmental growth factor that activates the ErbB class of tyrosine kinases receptors. It has a clear role in activating neurotransmitter receptors including the NMDARs [12]. BDNF is a neurotrophin predominantly present in the brain and is involved in neurogenesis and synaptic plasticity. Alterations in the levels of BDNF (both transcript and protein) observed in schizophrenia patients might lead to abnormal brain development and altered synaptic connectivity [13].

One of the major mechanisms for regulating NMDAR is mediated through microRNAs (miRNA) [14]. Studies also show that aberrations in post-transcriptional gene regulation, mediated by miRNAs, are associated with neurological disorders [15] Perkins et al in 2007 reported the first study on miRNAs in schizophrenia with custom microarray and qPCR using samples of human dorsolateral prefrontal cortex (DLPFC) and identified 15 differentially expressed miRNAs [16]. Similar studies were then carried out by other groups [17] in the superior temporal gyrus (STG) that showed a significant increase in the expression of several miRNAs. Understanding the association between miRNA expression and the function of its target gene will be crucial to understand disease mechanisms and to exploit miRNAs as biomarkers.

The aim of this paper is to understand the regulatory mechanisms of some of the miRNAs which might target NMDAR. We have identified putative miRNAs and validated some of them using in vitro culture systems. To validate the role of these miRNAs in vivo, we used a pharmacological model - MK-801 model of schizophrenia [18] and a neurodevelopmental model - the methylazoxymethanol acetate (MAM) model of schizophrenia [19].

## 2. Results

### 2.1. Computational prediction of miRNAs that bind to the 3’UTR regions of *Grin2A* and *Grin2B* transcripts

NMDARs are heterotetramers composed of multiple subunits, including two GluN1 subunits and two modulatory subunits from the GluN2 (GluN2A–GluN2D) and/or GluN3 (GluN3A, GluN3B) families [23]. To predict the miRNAs that regulate NMDAR we selected *Grin2A* and *Grin2B* (gene names of GluN2A and GluN2B sub-units respectively) as the targets because these are the major subunits of NMDAR that are involved in schizophrenia [24]. We used 3-5 online prediction tools for each target gene (Table 1), except miR-148b-3p, for which only two tools predicted interaction with *Grin2B*.

**Table 1.** MicroRNAs selected from different *in silico* prediction tools. Most of the selected miRNAs were predicted by three or more tools. miR-137 which is used as the positive control for *Grin2A* (Wright et al,2013) was also predicted in four tools. miR-223 which is used as the positive control for *Grin2B* (Harraz et al., 2012) was predicted only in two tools.

| <u>miRNAs for *Grin2A*</u> | <u>Different prediction tools which showed the selected miRNAs</u> |
| --- | --- |
| Rno-miR-148b (5p/3p) | microRNA.org, TargetScan, PITA |
| Rno-miR-296-3p | microRNA.org, MicroCosm, PITA |
| Rno-miR-129 (5p/3p) | microRNA.org, TargetScan, PITA |
| Rno-miR-137-3p | microRNA.org, TargetScan, PITA, MicroCosm |

**Table 1 .MicroRNAs selected from different *in silico* prediction tools.**
| <u>miRNAs for *Grin2B*</u> | <u>Different prediction tools which showed the selected miRNAs</u> |
| --- | --- |
| Rno-miR-185-5p | microRNA.org, miRDB, PITA, |
| Rno-miR-296-3p | microRNA.org, TargetScan, miRDB, PITA |
| Rno-miR-129 (5p/3p) | TargetScan, miRDB, PITA, MicroCosm |
| miR-148b (5p/3p) | TargetScan, microRNA.org |
| Rno-miR-223-5p | TargetScan, miRDB |

Additionally, we had used RNA hybrid tool to assess the stability of binding of the miRNA to the target 3’UTR. We obtained minimum 20-30 putative miRNAs per tool that targeted 3’UTRs of the NMDAR subunits. For further studies, we chose six miRNAs most of which were selected in a minimum of three prediction tools and were also reported to be differentially expressed in schizophrenia post mortem brain samples [16,17, 25, 26]. The target sites for these miRNAs were also conserved among vertebrates. Figure 1 shows the binding patterns of some of the miRNAs with the corresponding target along with the sequence alignment and conservation of the target. miR-148-5p and miR-129-2-3p were also predicted to bind with *Grin2A* (Suppl. Figure 1). Although miR-223-5p was shown to target *Grin2B* in only two tools in our analysis, it was already reported to regulate *Grin2B* and hence it was used as a positive control for *Grin2B* [27]. miR-137-3p, which was predicted by 3-4 tools, is reported to target and regulate *Grin2A* and hence this miRNA was taken as a positive control for *Grin2A* [28]. Thus, miRNAs such as miR-148b, miR-129-2 and miR-296 target both the receptor subunits whereas other miRNAs such as miR-185 target one of the subunits. Majority of them had a site type of 7mer-m8 or 7mer-A1 implying a strong interaction of miRNA seed sequence with the mRNA 3’UTR.

**Fig. 1.**
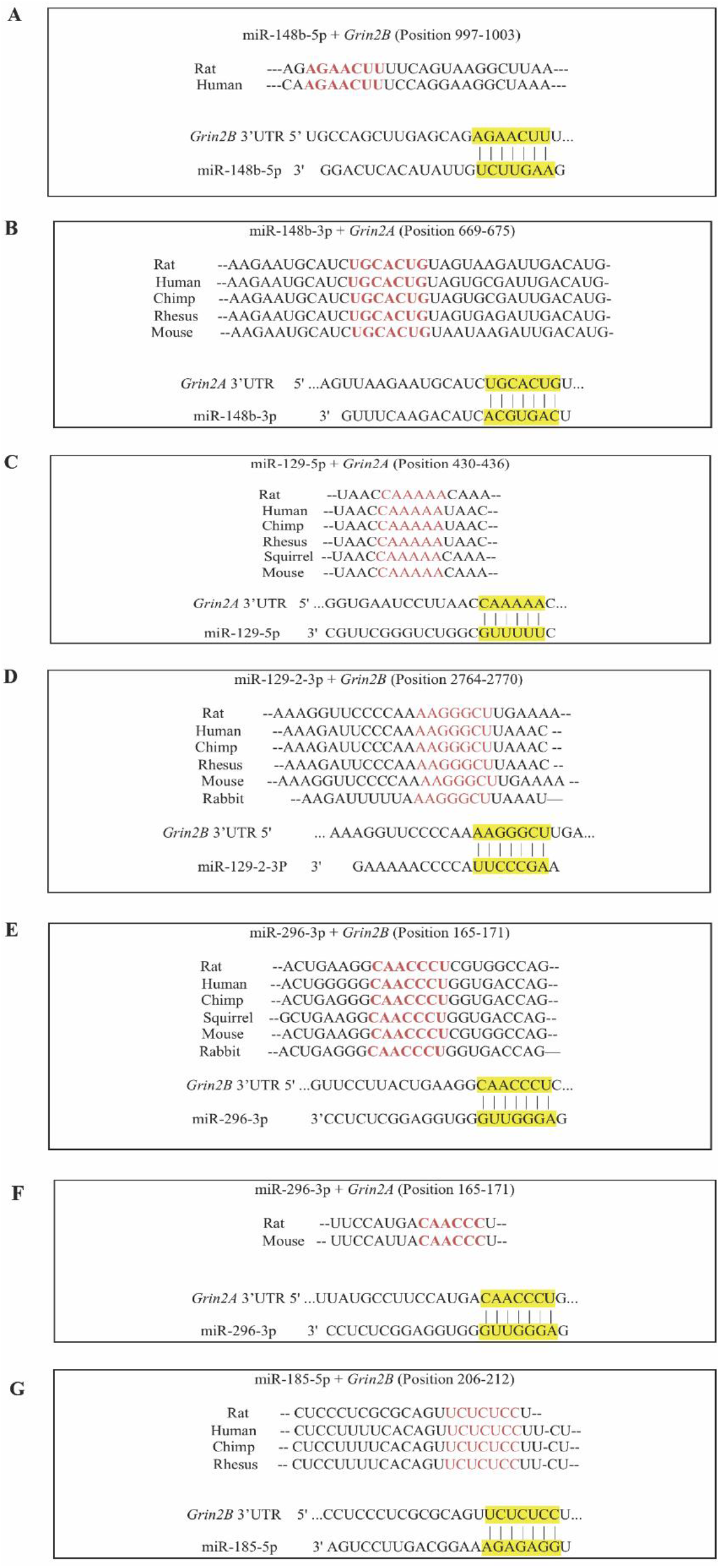
Sequence alignment of the binding site of miRNA in *Grin2B and Grin2A* showing conservation among different species. (A) Binding pattern of miR-148b-5p to *Grin2B* (B) Binding pattern of miR-148b-3p to *Grin2A* (C) Binding pattern of miR-129-5p to *Grin2A* (D) Binding pattern of miR-129-2-3p to *Grin2B* (E) Binding pattern of miR-296-3p to *Grin2B* (F) Binding pattern of miR-296-3p to *Grin2A* (G) Binding pattern of miR-185-5p to *Grin2B*

### 2.2. Experimental validation of the interaction between miRNAs and their targets

The miRNAs that were predicted to target the NMDAR subunits, were further screened for interaction with the corresponding 3’UTRs using the dual luciferase assay. The rat genomic sequences of the 3’UTRs of *Grin2A* and *Grin2B* and the pre-miRNA sequences were cloned into psiCHECK2 and pRIPM DsRed vectors respectively. Each pre-miRNA along with the corresponding 3’UTR were co-transfected into HEK-293 cells and were subjected to the assay. There was a significant decrease in the renilla luciferase activity for the precursor miRNAs-miR-129-2, miR-296, miR-223 and miR-148b against *Grin2B* 3’UTR (Figure 2A). There was no significant decrease in case of miR-185 implying weak or no interaction with the *Grin2B* 3’UTR. Similarly, four pre-miRNAs, miR-137, miR-129-2, miR-296 and miR-148b showed interaction with *Grin2A* 3’UTR (Figure 2B). These results implied that most of the predicted miRNAs are indeed interacting with the 3’UTRs of the *Grin2A* and *Grin2B* mRNAs in the cellular context.

**Fig 2.**
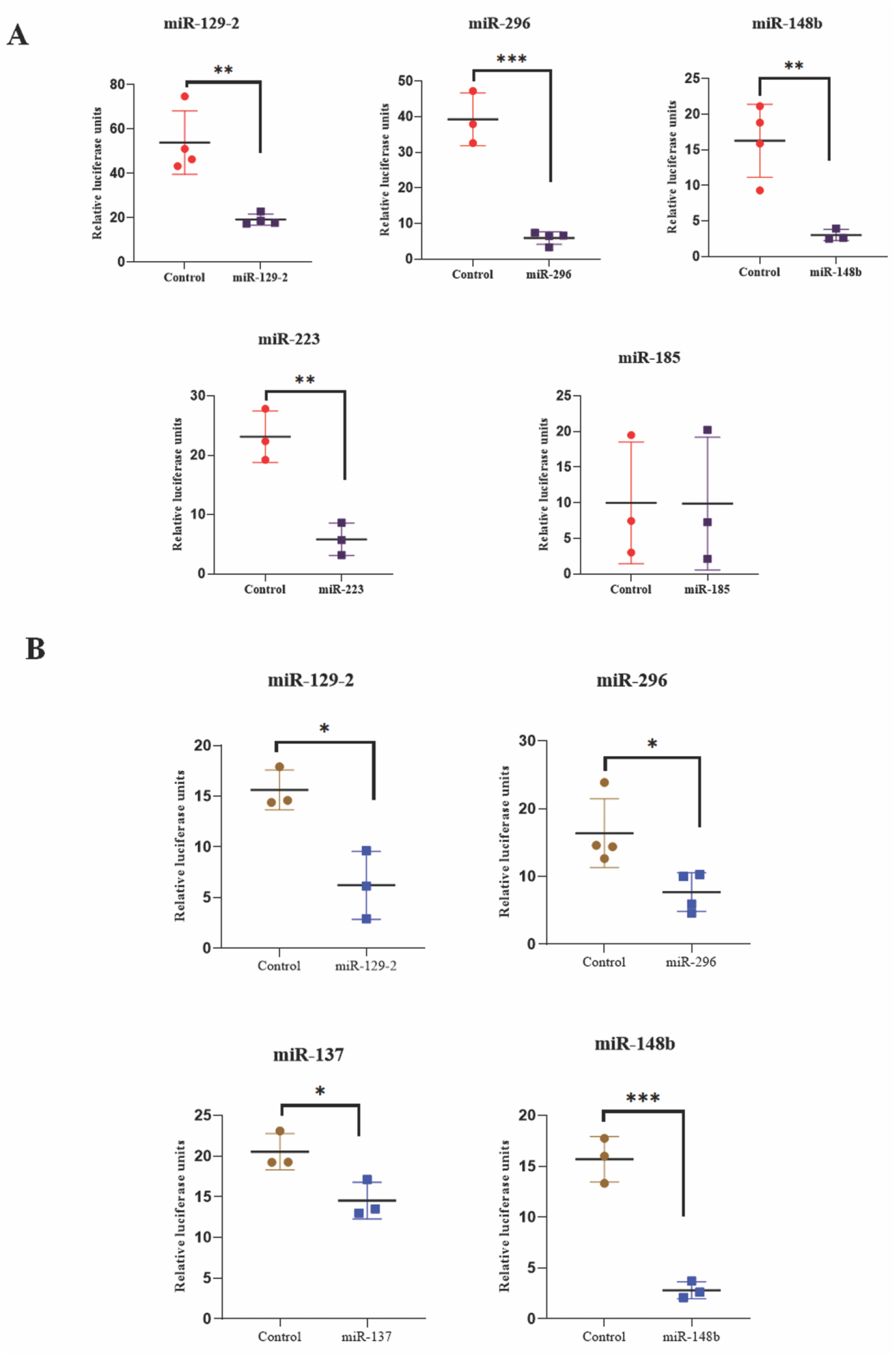
Quantitation of interaction between miRNAs and target sequences by dual luciferase assay. The dual luciferase assay was performed after co-transfecting the pre-miRNAs encoded in the pRIPM plasmid along with the psiCHECK2 plasmid carrying *Grin2B* 3’UTR (A) and psiCHECK2 *Grin2A* 3’UTR (B) plasmid to HEK-293 cell line. The control represents co-transfection of the pRIPM empty vector and psiCHECK2 *Grin2A/Grin2B* 3’UTR plasmids. The firefly luciferase activity arising from the psiCHECK2 3 ‘UTR vector is used for normalization. Statistical analysis was done by unpaired Student’s t test, ***p ≤ 0.001, **p ≤ 0.01, *p ≤ 0.05; n=3-4. Data are expressed as mean ±SD.

### 2.3. miRNAs downregulate the expression of GluN2B in primary neurons

To further support the results of the dual luciferase assay, the ability of the miRNAs in regulating the protein levels of NMDAR subunits in primary hippocampal neurons was tested. The temporal expression pattern of *Grin2B* transcripts were studied in primary hippocampal neurons maintained for different days *in vitro* (DIV 3 - DIV 14). It was found that *Grin2B* expression was extremely high in the initial days of culture (DIV 7-DIV 9) (Figure 3A). The miRNAs, miR-129-2, miR-296, miR-148b and miR-223, that were shown to interact with *Grin2B* in the bioinformatics analysis and dual luciferase assay, were transfected into hippocampal neurons at DIV 7 when the GluN2B expression was high. DsRed present in the pRIPM construct carrying the pre-miRNAs aided in marking the transfected neurons. The cells were fixed on DIV 10 and the expression of the GluN2B subunit was checked by immunocytochemical staining. GluN2B is majorly produced and processed in the cell body of the neurons and hence we checked for the somatic GluN2B expression [29]. The transfected cells showed significant reduction in GluN2B expression compared to the untransfected cells in the same microscopic field.

**Fig 3.**
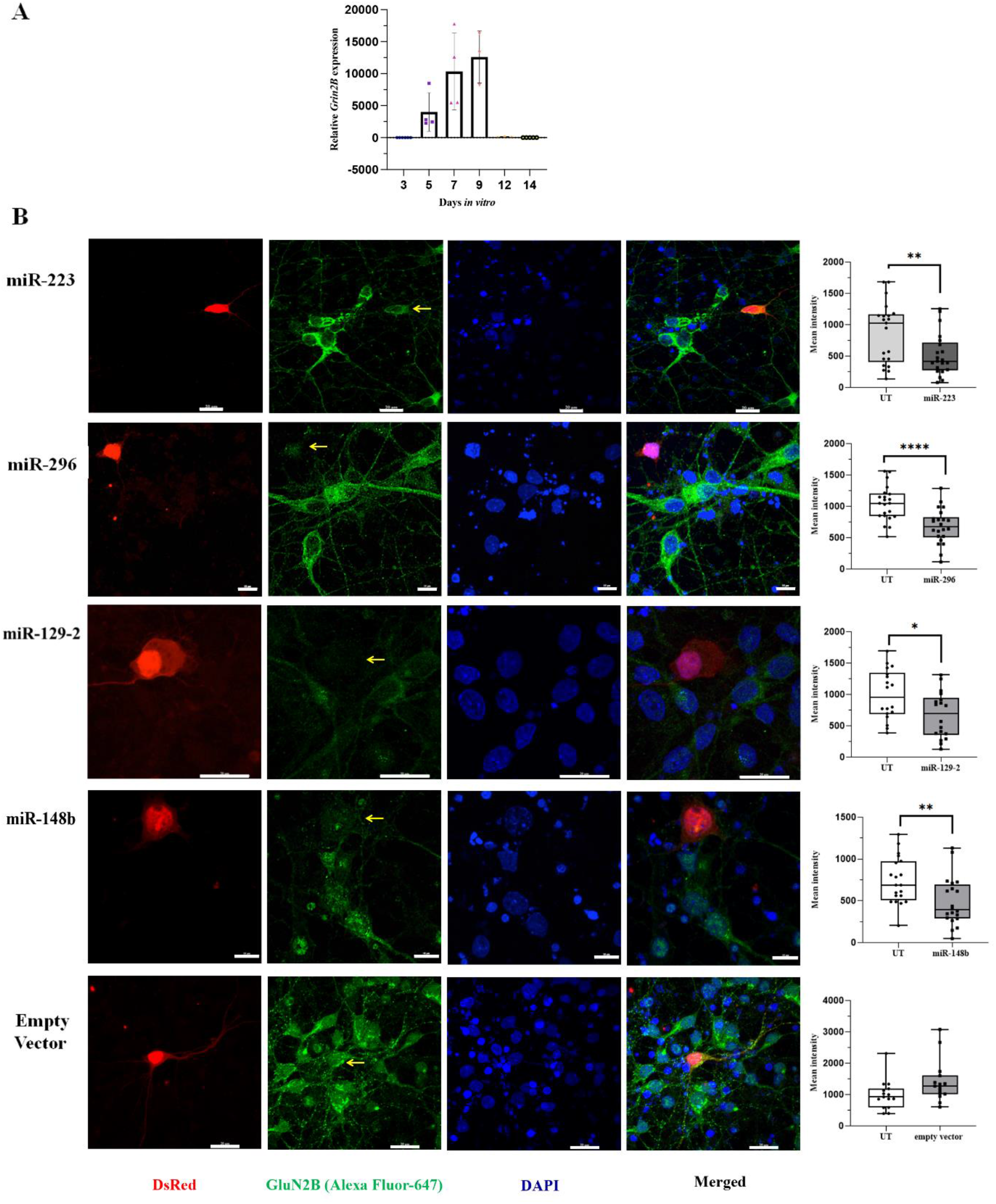
Evaluating the function of miRNAs in cultured neurons. (A) Temporal expression pattern of *Grin2B* transcript in primary neuronal cultures *in vitro*. The relative mRNA levels in rat hippocampal cultures from DIV3 to DIV14 are shown. The transcripts were quantified by SYBR qRT-PCR method and the values were normalized to beta actin. n=3-4. (B) Rat hippocampal neurons were transfected with pre-miRNAs encoded in the pRIPM vector carrying a DsRed reporter at DIV7. The cells were fixed at DIV 10 and were stained for GluN2B (with Alexa Fluor 455, green) and for the nucleus (with DAPI, blue). Images are representative of 3-4 experiments and were captured using Nikon confocal microscope. The transfected cells have the DsRed marker (red) and are indicated by arrows in the green images. Region of interest (ROI) quantification of fluorescence intensity for the untransfected (UT) and miRNA transfected cells were done using the NIS Elements V.4.0 imaging software. Most of fields had 1-2 transfected cells and 10 or more untransfected cells. For each field, the ROI quantification values were averaged for untransfected and transfected cells. Data are expressed as mean ±SD. Boxes indicate the lower quartile, median and upper quartile, and whiskers indicate the range of changes (minimum to maximum). Statistical analysis was done by unpaired Student’s t test: ****p ≤ 0.0001, ***p ≤ 0.001, **p ≤ 0.01, *p ≤ 0.05; n (Total no. of fields quantified in 3-4 experiments) = 15-22 (15-16 for Empty vector and 20-22 for the miRNAs). Scale bar: 10 μm for miR296 and miR148b; 20 μm for miR223, miR129-2 and empty vector.

The empty pRIPM vector was used as a control, which showed no observable change in GluN2B expression in the transfected cells compared to untransfected cells (Figure 3B). By performing immunocytochemical analysis, it has been possible to visually appreciate the decrease in expression in somatic GluN2B in the miRNA-overexpressed neurons. Our aim was to show that the protein level was brought down by the predicted miRNAs, which was achieved by studying GluN2B. Overall, these results signify that the miRNAs that showed interaction with the target 3’UTR in the dual luciferase assay, also caused downregulation in protein expression in cultured neurons for GluN2B. It is highly probable that these predicted miRNAs might also downregulate GluN2A expression levels.

### 2.4. Regulation of NMDAR by the selected miRNAs in vivo - Pharmacological and neurodevelopmental models of schizophrenia

The data so far shows that the selected miRNAs can interact with the 3’UTRs of *Grin2A* and *Grin2B*. Regulation of protein expression by the miRNAs could also be shown using GluN2B in primary neurons as the model. To understand the role of these miRNAs in schizophrenia, we used two rat models of the disease, the MK-801 (pharmacologi-cal) model and the MAM (neurodevelopmental) model. Since these miRNAs and the NMDAR subunits are known to undergo altered regulation in schizophrenia in humans [4,5,17] their levels were monitored in the animal models towards investigating whether alterations in the regulation of the NMDAR subunits by miRNAs could contribute to the disease.

#### 2.4.1. Treatment regimes for the animal models

MK-801 is a non-competitive NMDAR channel blocker. It was intraperitoneally administered in two different doses - 0.5 mg/Kg and 1.0 mg/Kg for five days based on previous reports [30–33]. The resulting blockage of NMDAR is expected to cause alterations in the brain similar to schizophrenia. Following this a five-day drug washout period was allowed so that the observed behavior reflects the disease condition and not the effect of the drug *per se* (Figure 4A).

**Fig 4.**
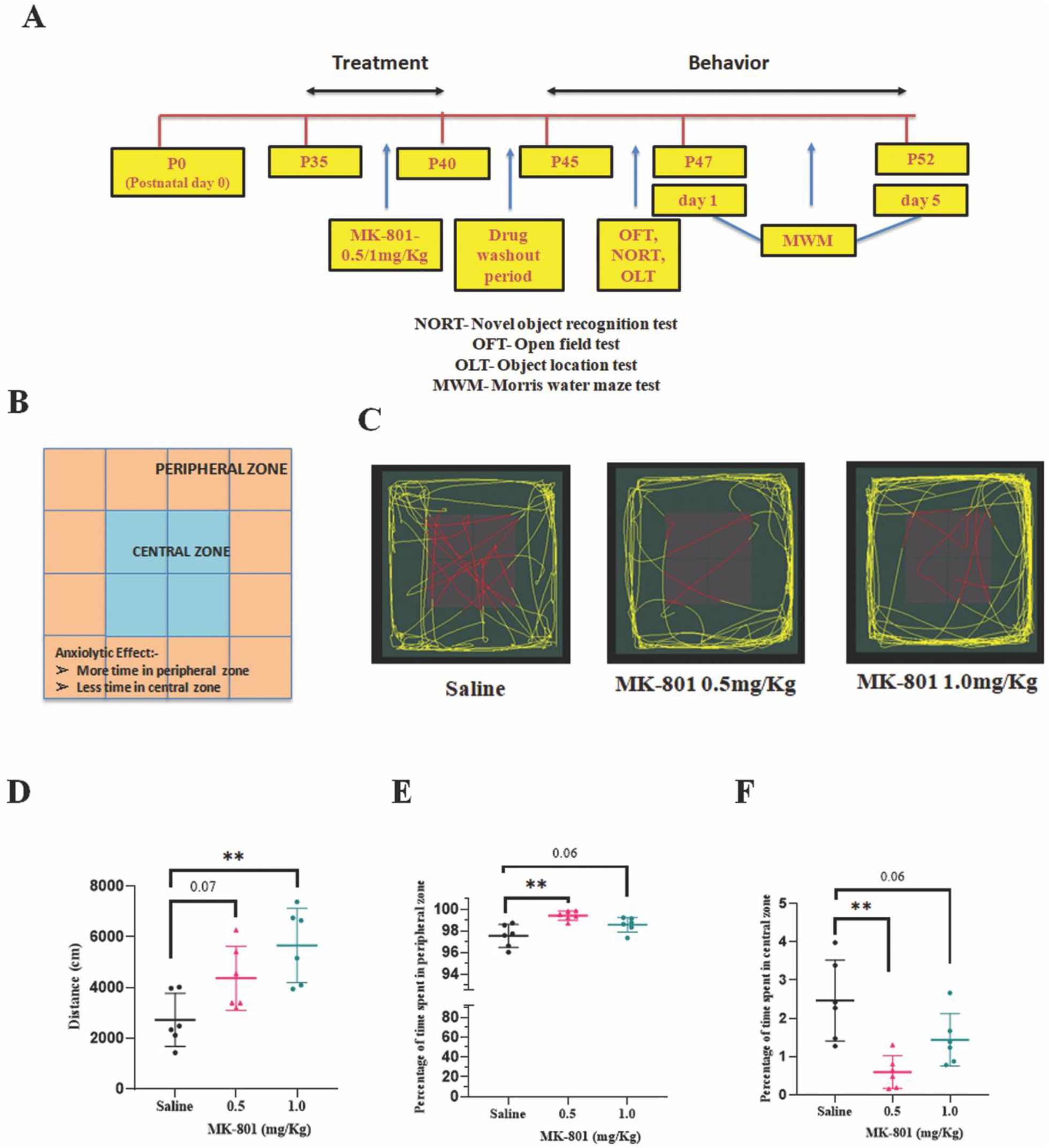
MK-801 treated animals show anxiolytic behavior. (A) Timeline of experimental procedures for MK-801 model. (B) The schematic of the box used for the open field test. (C) Representative tracks of movement of the animals inside the box during 10-11 min. (D) Total distance travelled by the animals during the open field test. (E) Time spent in the peripheral zone and (F) central zone during the total period of 10-11 min. Quantitation is shown as mean ± SD; p values obtained by one-way ANOVA followed by Dunnett’s multiple comparison test are shown, **p ≤ 0.01, n=6 per group.

MAM is a neurotoxin that affects neurogenesis in the central nervous system of embryos when injected on gestational day (GD) 17. The pups of MAM treated dams have been shown to exhibit structural and functional alterations observed in schizophrenia [19, 34]. MAM was administered intraperitoneally in the dams at GD 17 stage. The pups were weaned at postnatal day 30 (P30) and were used for the behavioral experiments from P45 onwards (Figure 6A).

**Fig 5.**
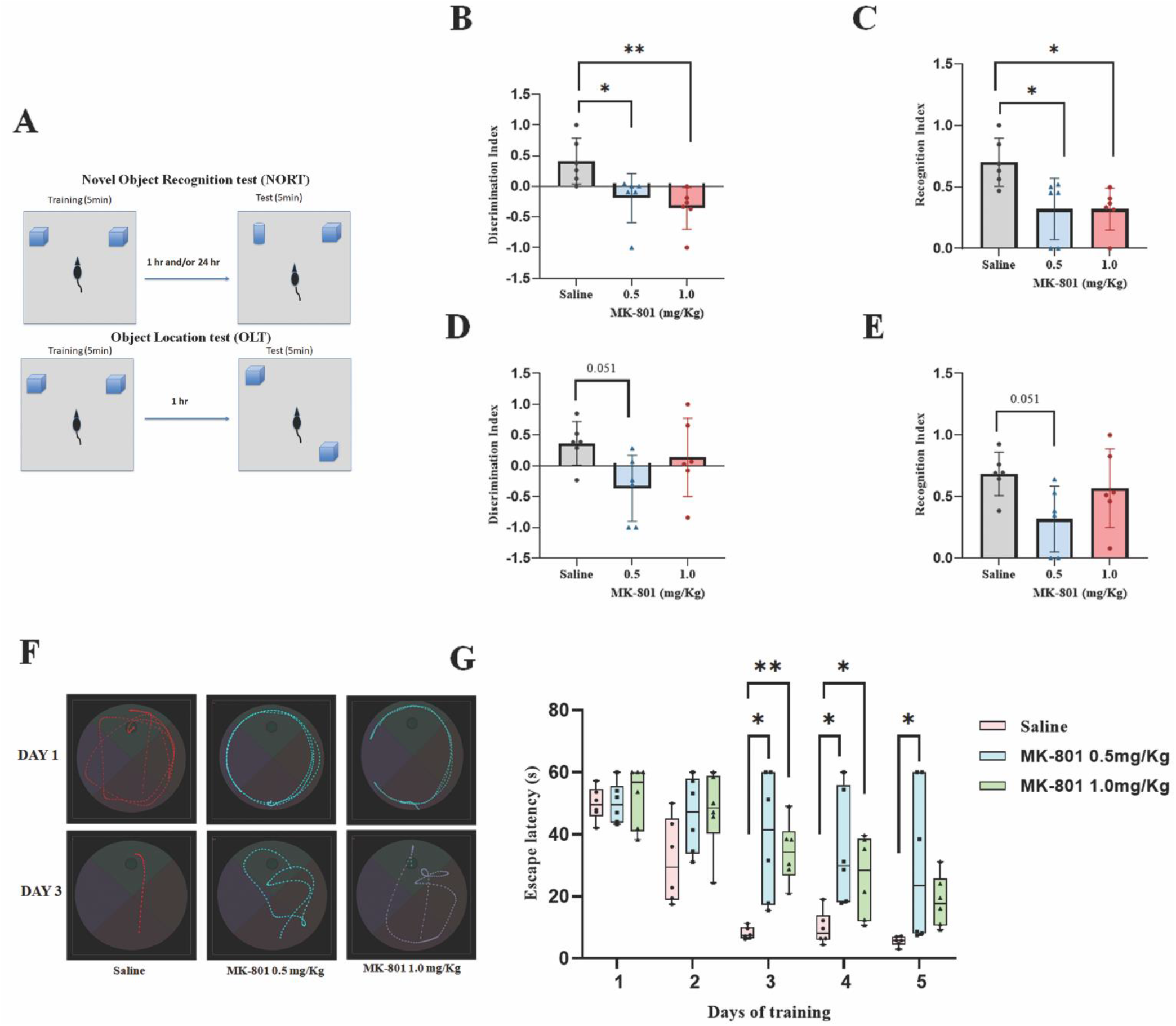
MK-801 treated animals show impaired learning and memory. (A) Schematics representation of NORT/OLT. (B) DI and (C) RI in NORT. (D) DI and (E) RI in OLT. Quantitation is shown as mean ±SD. Statistical analysis was done by one-way ANOVA followed by Dunnett’s post hoc multiple comparison test with p values indicated: **p ≤ 0.01, *p ≤ 0.05; n=6 per group. (F) Representative swim tracks of saline treated and MK-801 treated animals in MWM on day 1 and day 3 of training. (G) Quantitation represented as mean ±SD of the latency to reach the platform of the experimental groups. Boxes indicate the lower quartile, median and upper quartile, and whiskers indicate the range of changes (minimum to maximum). **p ≤ 0.01, *p ≤ 0.05, by repeated measures two-way ANOVA with Dunnett’s multiple comparison test; n=6 per group.

#### 2.4.2. Behavioral test reveals schizophrenia-like phenotype

##### 2.4.2.1. OFT

Initially, the animals were subjected to OFT. This test is used for analyzing anxiety levels and locomotor ability. It was observed that the MK-801 treated rats were hyperactive as seen by the significantly longer distance travelled by them compared to the controls (Figure 4C and D). Analysis of the data by one-way ANOVA test indicated significant differences among the three groups (F (2,15) = 8.03; p=0.0042). The p values for pairwise comparison by the Dunnett’s test is shown in Figure 4D. It was also observed that the MK-801 treated rats (0.5 mg/Kg and 1.0 mg/Kg) spent more time in the peripheral zone and less time in the central zone of the box indicating anxious behavior (Figure 4E and 4F) [35]. The percentage of time spent in the peripheral and central zone showed significant differences among the groups (One way ANOVA: F (2,15) = 8.93; p=0.0028).

The MAM treated animals covered significantly lesser distance compared to the control group (Student’s unpaired t test: t=3.05, p=0.0066) (Figure 6B and 6C) [35]. The MAM treated rats also spent more time in the peripheral zone and less time in the central zone indicating anxiety (Student’s unpaired t test: t=2.076, p=0.051 for both the parameters) (Figure 6D and 6E).

##### 2.4.2.2. NORT

We conducted tests for learning and memory such as NORT and OLT. It was observed that the recognition index (RI) and the discrimination index (DI) in NORT for the MK-801 treated (both the doses) animals were significantly less compared to their respective saline treatment group, which indicates that the treated animals spent less time in exploring the novel object (Figure 5B, 5C). The one way ANOVA test indicated significant differences among the groups for DI (F(2,15) = 7.10; p=0.0067) and RI (F(2,15) = 6.68; p=0.0084).

The RI and DI in NORT for the MAM treated animals were lesser compared to their respective saline treatment group, which indicates that the treated animals did not spend more time in exploring the novel object (Figure 6F and 6G). The data for the MAM animals was analyzed using the unpaired Student’s t test. The MAM treated animals showed learning deficit in the NORT conducted at both 1 hr (t=1.744, p=0.095 for DI; t=2.24, p=0.035 for RI) (Figure 6F and 6G) and 24 hrs (t=4.14, p=0.0005 for DI; t=4.39, p=0.0003 for RI) (Figure 6H and 6I) after training [36].

##### 2.4.2.3. OLT

The RI and DI of MK-801 treated animals in OLT were also lesser compared to their respective controls implying that the treated animals did not spend more time with the displaced object compared to the stationary/non-displaced object (Figure 5D and 5E). The ANOVA analysis did not show significant difference among groups (F(2,15) = 3.077; p=0.0759 for DI and RI). However, the Dunnett’s analysis showed that the 0.5 mg/Kg treated animals could be having a defect in DI and RI (p=0.051).

There was major difference between the saline group and the MAM treated group as indicated by RI and DI (t=5.08, p<0.0001 for DI; t=5.411, p=0.0001 for RI) in OLT (Figure 6J and 6K). These results showed that the MK-801 and the MAM treated animals have cognitive deficits showing impairment in learning and memory.

##### 2.4.2.4. MWM test

We also conducted MWM test to study the hippocampus dependent learning and memory phenotype for these animals. The treated animals of both models took significantly more time to reach the platform on day 3 when compared to saline group. The MK-801 treated animals showed improvement by day 4 and 5 but the time taken to reach the platform was still significantly higher compared to the control group indicating that the treated rats have working memory deficit (Figure 5F and 5G). The repeated measures two-way ANOVA indicated significant differences be-tween the time and groups of the MK-801 cohort (F(8,60) = 2.87; p=0.0088).

The MAM treated animals did not show learning behavior till day 3 (Figure 6L and 6M). By day 5, they showed minimal learning when compared to their performance on day 3 although still impaired compared to the control group (Figure 6M). The repeated measures two-way ANOVA indicated significant differences between the time and groups (F(4,84) = 9.72; p<0.0001).

**Fig 6.**
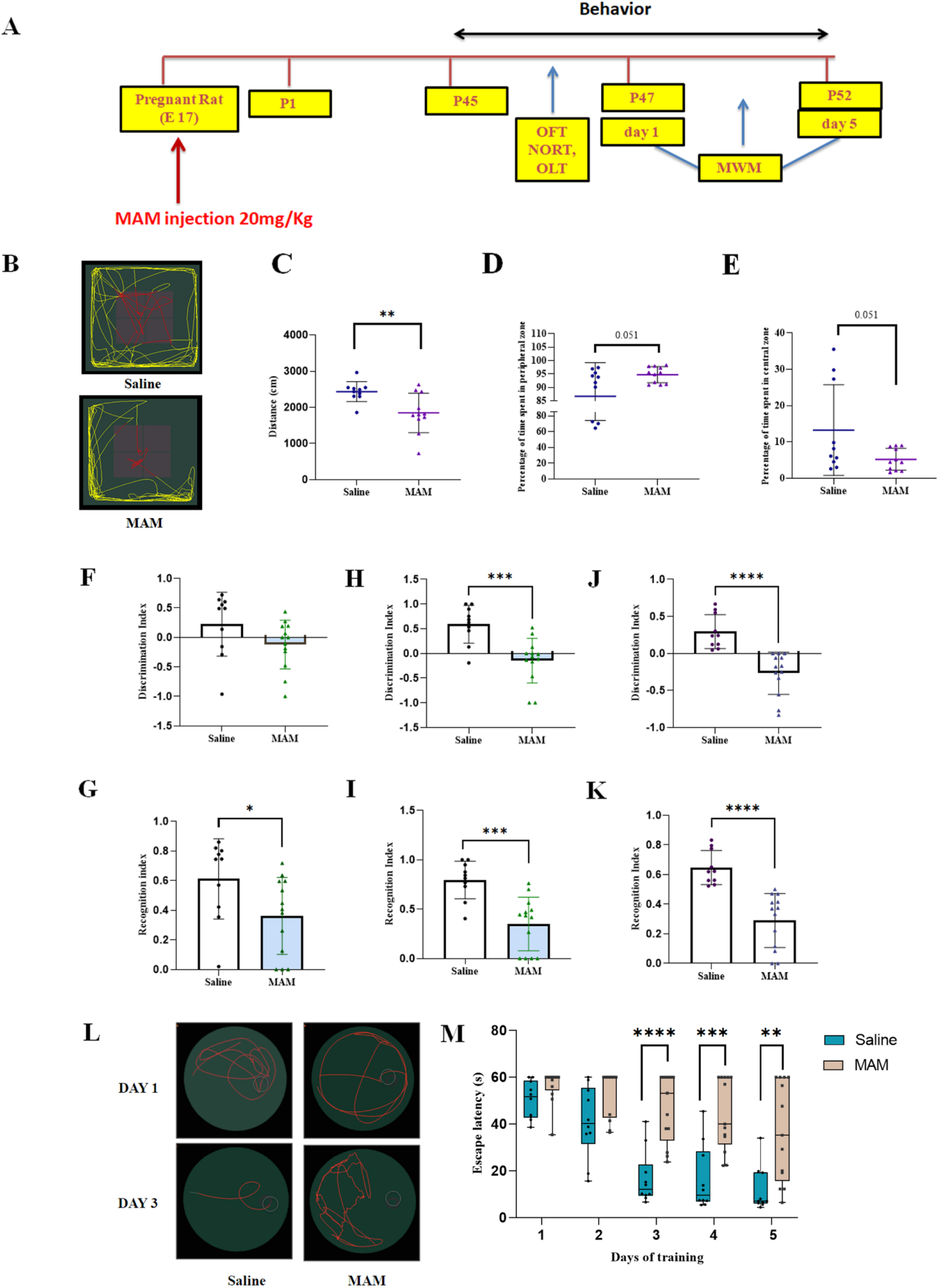
MAM treated animals show decline in cognitive function. (A) Timeline of the experimental procedures followed for MAM model of schizophrenia. (B) Representative tracks of movement of the animals during 10-11 min when placed inside the box in OFT. (C) Total distance travelled by the animals during OFT. (D) Time spent in the peripheral zone and (E) central zone during the total period of 11min. (F and G) DI and RI in NORT conducted 1hr after training. (H) DI and (I) RI in NORT conducted 24 hrs after training. (J) DI and (K) RI in OLT conducted 1hr after training. (L) Representative swim tracks of saline injected and MAM injected animals in MWM test on day 1 and day 3 of training. (M) Quantitation represented as mean ±SD of the latency to reach the platform of the two groups. Boxes indicate the lower quartile, median and upper quartile, and whiskers indicate the range of changes (minimum to maximum). Statistical analysis for OFT, NORT and OLT was done by Student’s t test and the p values are indicated: ****p ≤ 0.0001,***p ≤ 0.001, **p ≤ 0.01, *p ≤ 0.05; n=10-13 per group. Statistical analysis for MWM test was done through repeated measures two-way ANOVA with Sidak’s test: ****p ≤ 0.0001, ***p ≤ 0.001, **p ≤ 0.01, n=10-13 per group.

#### 2.4.3. Molecular changes in the hippocampus of the MK-801 and MAM models of schizophrenia

##### 2.4.3.1. Alterations in protein levels

We checked the protein levels of the NMDA receptor subunits, GluN2A and GluN2B in the hippocampal region of both models and found that they were significantly reduced compared to the saline group (Figure 7A, B, F and G). One way ANOVA revealed significant effects among the MK-801 groups (F(2,12) = 5.76, p=0.017 for GluN2A; F(2,11) = 5.47, p=0.022 for GluN2B). Dunnett’s post hoc comparison tests indicated significant decreases in the protein expression of GluN2A and GluN2B for both the MK-801 treatment groups compared to the saline group. We al-so checked the levels of other proteins implicated in schizophrenia such as neuregulin 1 (NRG1), CaMKIIα and brain derived neurotropic factor (BDNF) and found that all of these were upregulated in the animals treated with 1.0 mg/Kg of MK-801 (Figure 7C, D and E). Their ANOVA analysis results were F (2,9) =4.92, p=0.03 for NRG1, F(2,9) =4.54, p=0.04 for BDNF and F(2,10) =5.08, p=0.03 for CaMKIIα.

**Fig 7.**
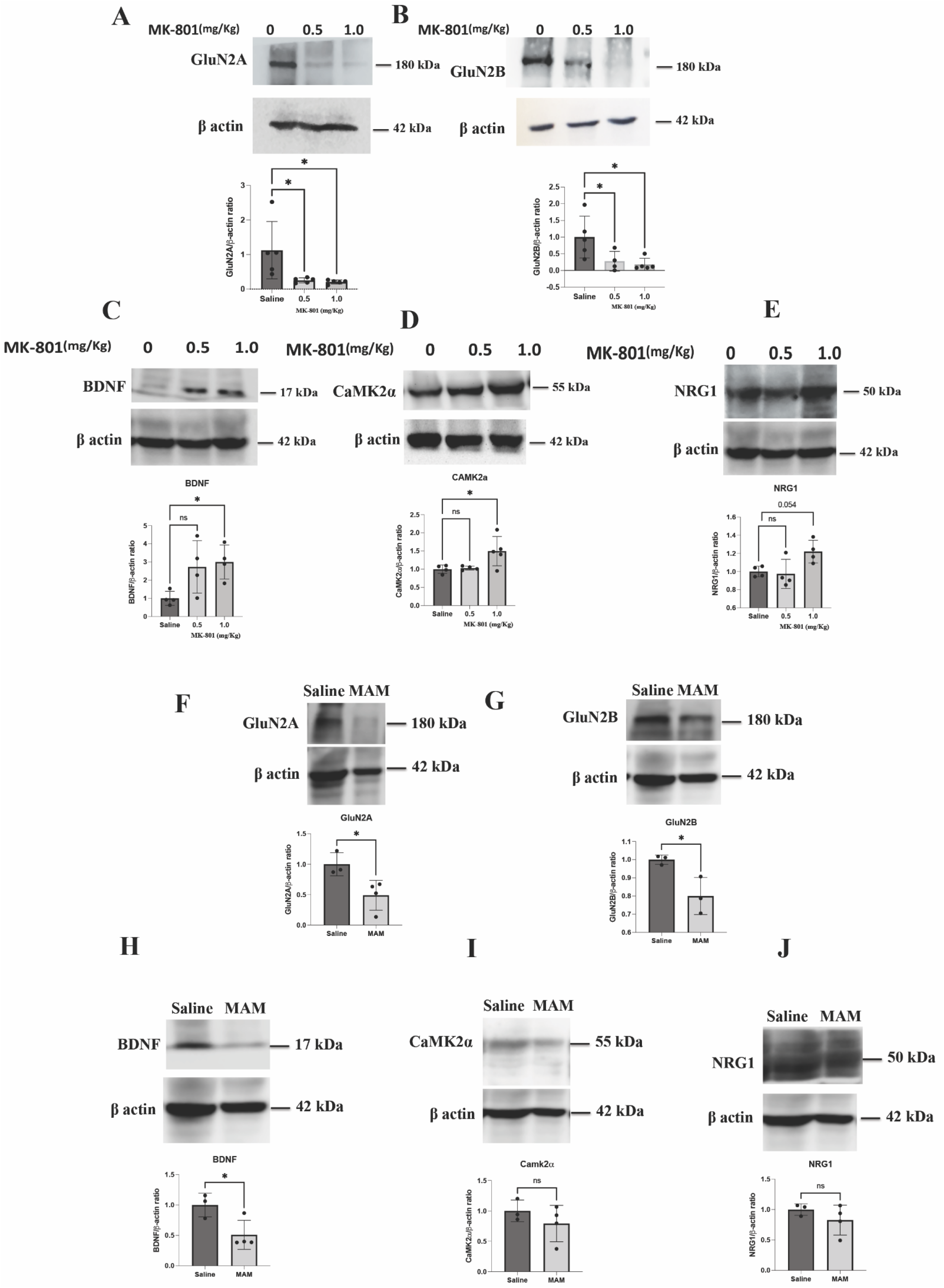
Changes in the protein expression level in the hippocampi of the MK-801 and MAM models of schizophrenia. (A-E) Representative western blots of GluN2A, GluN2B, BDNF, CaMKIIα and NRG1 for the MK-801 model are shown. n=4-5 (F-J) Representative western blots of GluN2A, GluN2B, BDNF, CaMKIIα and NRG1 of MAM model are shown. n=3-4 Mean ±SD of quantification of the band intensity of the protein normalized to band intensity of actin for multiple samples are also shown. Statistical analysis was done by Student’s t test for MAM group and one-way ANOVA with Dunnett’s test for MK-801 group. **p ≤ 0.01, *p ≤ 0.05.

For the MAM model also, we found significant decreases in protein levels of GluN2A (t=2.96, p=0.03) and GluN2B (t=3.30, p=0.02). However, the changes in the levels of BDNF, CaMKIIα and NRG1 were different compared to the MK-801 model (Figure 7H, I and J). There was a significant reduction in the BDNF levels (t=2.89, p=0.03) but no significant change was observed for NRG1 (t=1.13, p=0.30) and CaMKIIα (t=1.06, p=0.33).

##### 2.4.3.2. Alterations in mRNA levels

The transcript levels of these proteins were subsequently checked. Transcript of *Grin2A* was found to be significantly upregulated in the hippocampal region in MK-801 model of schizophrenia (Figure 8A). The one-way ANOVA analysis showed significant differences between the groups for *Grin2A* (F (2,10) =13.8; p=0.001) and *Grin2B* (F (2,11) =4.87; p=0.03) with Dunnetts’s post hoc tests showing significant up-regulation of *Grin2A* and *Grin2B* in both the doses of MK-801 (Figure 8 A). The CaMKIIα transcript was observed to be significantly upregulated in the 0.5 mg/Kg dose of MK-801 treatment using Dunnett’s test and overall significance was observed among the groups using the one-way ANOVA test (F (2,9) =4.49; p=0.04). Significant differences were not found with one way-ANOVA for *NRG1* and *BDNF* transcript levels among the groups of MK-801 treatment (F (2,11) =3.23; p=0.08 for *NRG1*, F (2,12) =3.44; p=0.06 for *BDNF*). Upregulation in the *BDNF* (significant) and *NRG1* (p=0.0507) levels with respect to saline group upon treatment with 1.0 mg/Kg of MK-801 group was observed with Dunnett’s post-hoc test. (Figure 8 A).

**Fig 8.**
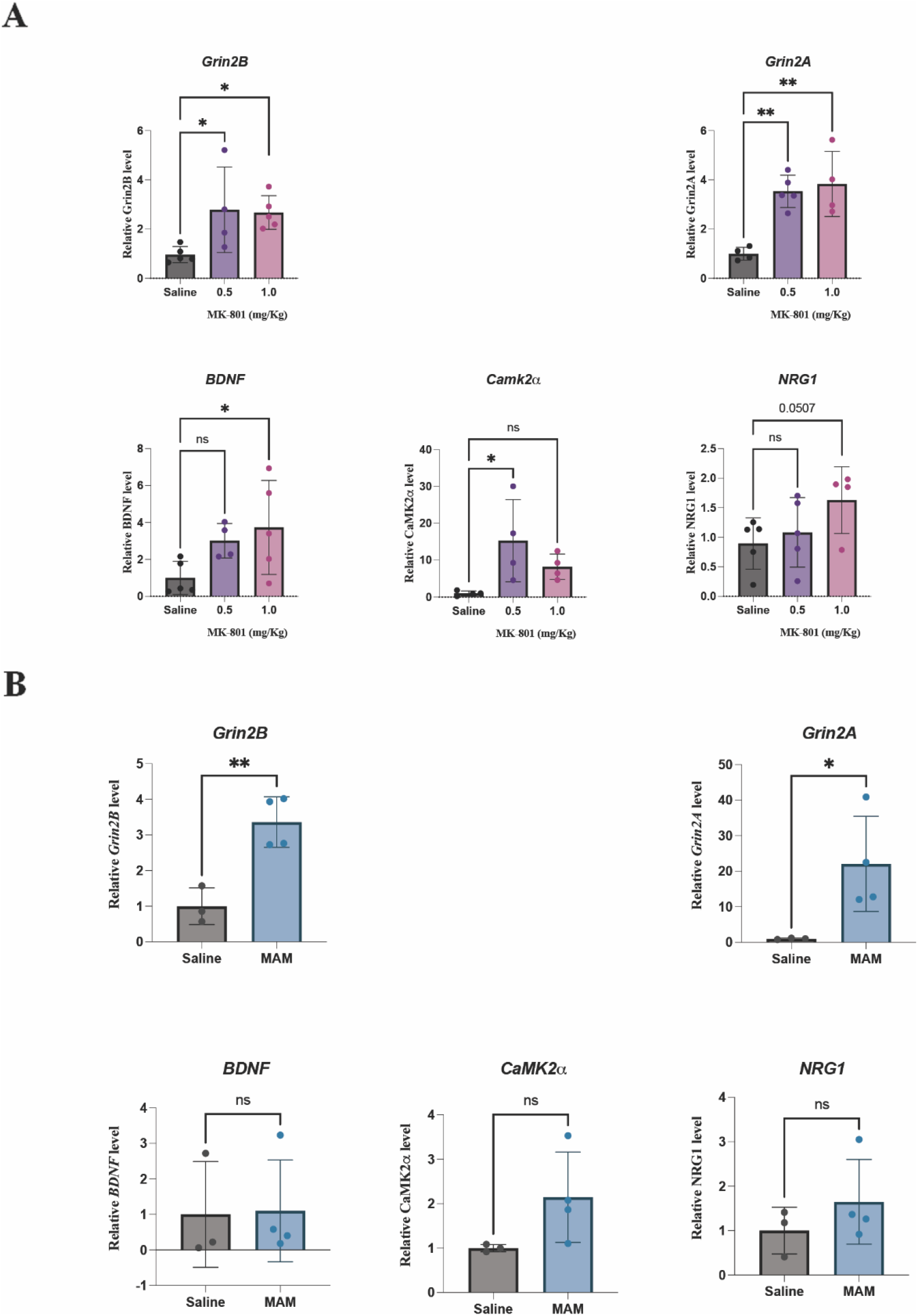
Changes in mRNAs levels in the hippocampal region. Quantitation by qPCR, represented as mean ±SD of relative expression of *Grin2A*, *Grin2B*, *BDNF*, *CaMKIIα and NRG1* in the hippocampal region of the MK-801 model (A) and MAM model (B) is shown. Statistical analysis was by one-way ANOVA with Dunnett’s test for MK-801 group and Student’s t test for MAM group:**p ≤ 0.01,*p ≤ 0.05; n=4-5 for MK-801 group, n=3-4 for MAM group.

There was a significant increase in the *Grin2A* and *Grin2B* transcript levels in MAM model (t=2.65, p=0.04 for *Grin2A;* t=4.83, p=0.005 for *Grin2B* by Student’s t test analysis) (Figure 8B). The transcript levels of *CaMKIIα* were upregulated but not significantly (t=1.90, p=0.11) and the transcripts of *NRG1* and *BDNF* were not significantly affected in the MAM model (t=1.04, p=0.34 for *NRG1*; t=0.09, p=0.93 for *BDNF*) (Figure 8B).

##### 2.4.3.3. Alterations in miRNA levels

We next measured the levels of the selected miRNAs, which are predicted to target the NMDAR subunits, in the hippocampal tissues of the animal models. It was observed that some of these miRNAs were significantly upregulated in the treated groups. The miRNAs, miR-296-3p and miR-148b-3p were significantly upregulated in both the doses of MK-801 treatment (Figure 9A). The one-way ANOVA indicated significant difference among the groups for both these miRNAs (F (2,12) =8.76; p=0.004 for miR-296-3p and F (2,11) = 5.60; p=0.02 for miR-148b-3p). The one-way ANOVA for miR-223-5p and miR-129-2-3p also showed significant differences between the groups (F (2,11) =6.46; p=0.013 for miR-129-2-3p, F (2,12) =4.33; p=0.038 for miR-223-5p) with Dunnett’s multiple comparison test showing significant upregulation in the 0.5 mg/Kg MK-801 treatment group for both the miRNAs. miR-137-3p was significantly upregulated upon 1 mg/Kg MK-801 treatment with ANOVA test showing significant difference among the groups (F (2,11) =8.08; p=0.006) (Figure 9A).

**Fig 9.**
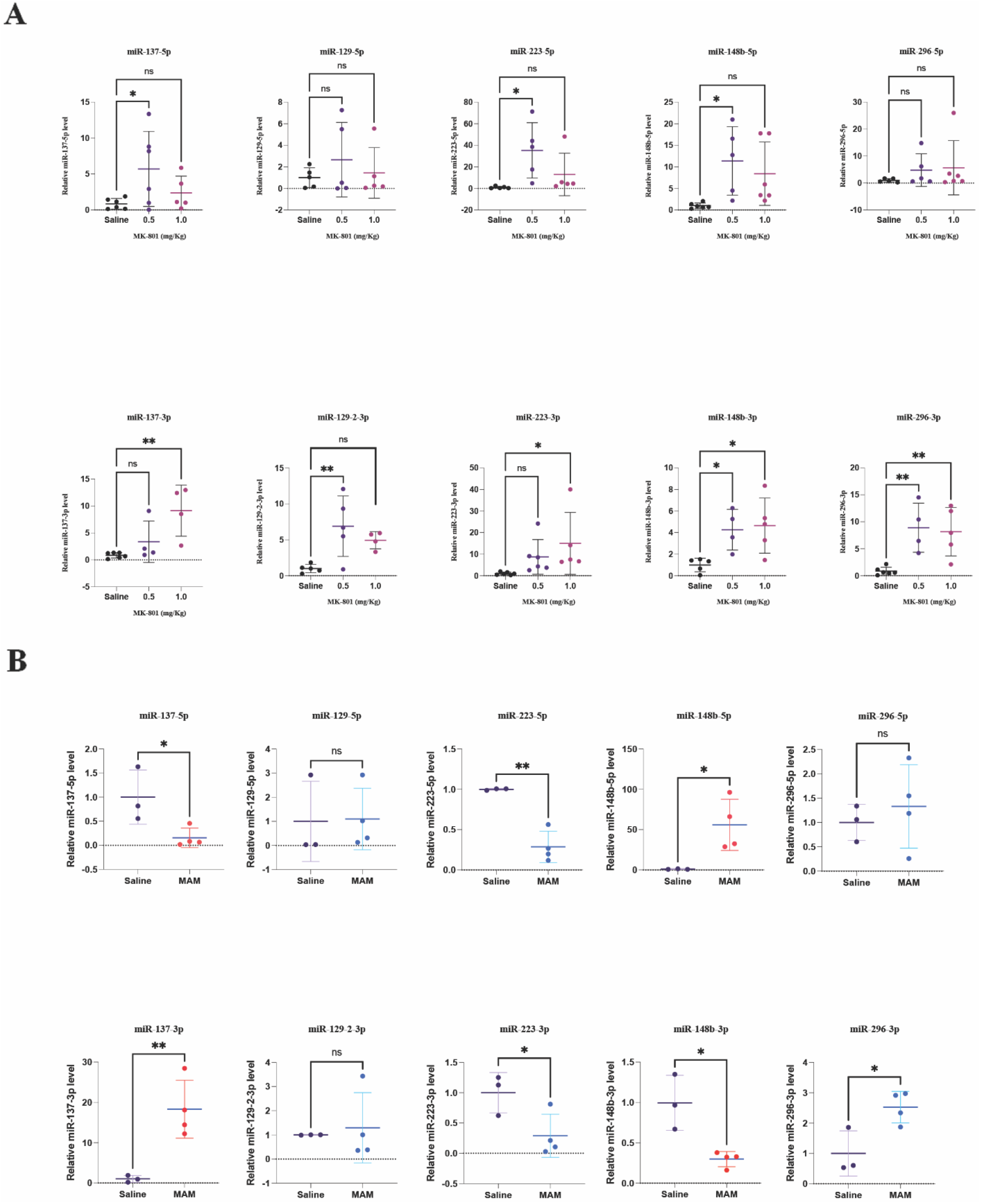
Alterations in the miRNA levels in schizophrenia animal models. Quantitation by qPCR, represented as mean ±SD, of the relative levels of −5p and-3p forms of miRNAs in the hippocampal region of the MK-801 model (A) and MAM model (B). Statistical analysis was done by one-way ANOVA with Dunnett’s test for MK-801 group and Student’s t test for MAM group. ****p ≤ 0.0001, ***p ≤ 0.001, **p ≤ 0.01, *p ≤ 0.05. n=4-6 for MK-801 group; n=3-4 for MAM group.

We also estimated the levels of the complimentary strands of these miRNAs and found that miR-137-5p (for MK-801, 0.5 mg/Kg dose), miR-223-3p (for MK-801, 0.5 mg/Kg dose) and miR-148b-3p (for MK-801, 1.0 mg/Kg dose) showed significantly high levels in the MK-801 model. The one-way ANOVA test indicated significant differences among the groups for only miR-148b-5p (F (2,14) =4.30; p=0.03 for miR-148b-5p; F (2,14) = 3.19; p=0.07 for miR-137-5p, F (2,14) = 3.28 and p=0.06 for miR-223-3p). The changes in the levels of miR-129-5p and miR-296-5p were not significant when analyzed by one-way ANOVA test (F (2,12) = 0.61; p=0.55 f or miR-129-5p, F (2,13) = 0.64; p=0.53 for miR-296-5p) or by Dunnett’s test (Figure 9A).

We measured these miRNAs in the MAM treatment and observed a similar pattern of upregulation in the MAM model for miRNAs miR-137-3p, miR-296-3p and miR-148b-5p (t=4.04, p=0.009 for miR-137-3p; t=3.21, p=0.02 for miR-296-3p; t=2.92, p=0.03 for miR-148b-5p, by t test) (Figure 9B). However, miR-129-2 did not show any significant change in the MAM model (t=0.34, p=0.74 for miR-129-2-3p; t=0.08, p=0.90 for miR-129-5p, by t test), suggesting that it might not have a role in this condition. We also noticed a pattern of inverse responses by the 3p and 5p forms of miR-148b and miR-137 in the MAM model. While the miR-148b-3p and miR-137-5p forms were significantly decreased in the MAM model (t=4.0, p=0.01 for miR-148b-3p; t=2.85, p=0.03 for miR-137-5p, by t test), their complimentary strands, miR-148b-5p and miR-137-3p showed significant increase. Both forms of miR-223 were significantly decreased (t=2.68, p=0.04 for miR-223-3p; t=6.20, p=0.01 for miR-223-5p) in the MAM model implying that it might not be regulating NMDAR in this model (Figure 9B).

The miRNAs that have elevated levels cause translational arrest of the upregulated NMDAR subunit transcripts thereby causing the observed reduction in GluN2A and GluN2B protein levels. This data suggests that miR-148b-5p, miR-296-3p and miR-129-2-3p might be strong candidates in regulating NMDAR *in vivo* apart from miR-137 and miR-223, which have already been shown to regulate the GluN2A and GluN2B subunits respectively.

## 3. Discussion

Altered expression pattern of miRNAs in neuropsychiatric disorders like schizophrenia and autism as well as in neurodegenerative disorders such as Alzheimer’s and Huntington’s diseases has been reported [37] but whether and how miRNA dysregulation leads to such disease phenotypes is still unclear. Our results demonstrate that one of the mechanisms of action of miRNAs miR-148b, miR-129-2 and miR-296 in schizophrenia, might be through downregulating the subunits of NMDAR. The use of multiple prediction tools with different algorithms helped to reduce false positives. Most of the miRNAs selected after computational prediction showed interaction with their respective target in the luciferase assay as well as in cultured primary hippocampal neurons. However, it was not possible to determine which strand of the miRNAs, 5p or 3p is actually targeting the protein using the transfection-based assays as the plasmid contains the precursor form which has both the arms of the miRNA.

The MK-801 model represents general NMDAR hypofunction and the MAM model recapitulates schizophrenia with developmental etiology [19,38]. The phenotype of the MK-801 model has been correlated to the positive and cognitive symptoms of schizophrenia [39]. MAM causes developmental anomalies leading to schizophrenia-like symptoms such as cognitive dysfunction, reduced social interaction and deficits in sensorimotor gating [19,40]. It is known that schizophrenia majorly occurs in men of 20-30 yrs of age and hence we chose male Wistar rats at P40-45 days of age that represents adolescence to early adulthood for the MK-801 model [41–44]. We did not use female Wistar rats for the MK-801 model, which is a limitation to our study. Female animals were included in the MAM model along with the males as there are very few reports on MAM models of schizophrenia using female animals [45,46]. Although the doses of 0.5 mg/Kg and 1.0 mg/Kg of MK-801 were used previously in separate studies, our study involves detailed comparative analysis of behaviour upon administration of these doses [29–32, 47–49]. In our treatment regime, the washout period of 5 days ensures that the drug *per se* won’t interfere with the behavior and hence the observed behavioral changes could be arising from the drug-induced changes in the brain tissue (Figure 4 and 5). It is also to be noted that the animals showed molecular changes in the brain tissue even when they were sacrificed 15-20 days after the MK-801 injections.

The behavioral parameters observed for the MAM model were more or less similar to those of the MK-801 model reflecting deficits, except that the MAM-treated animals were hypoactive (Figure 6B, C, D and E) whereas the MK-801 treated animals showed hyperactivity (Fig 4C, D, E and F). Both these are indicative of emotional imbalances known to occur in schizophrenia [50]. Cognitive defects revealed in the NORT, OLT and MWM test occurred in both the models (Figure 5 and Figure 6 F-M). We did not find any significant differences between the performance levels of the males and the females of the MAM model implying that this model might not have any gender bias under our conditions (data not shown). Since performance of these tasks involves the hippocampal region [51,52], the results indicated alterations in the hippocampal region, most likely with the proteins involved in synaptic plasticity.

Consistent with observed impairments in synaptic plasticity, the protein levels of GluN2B and GluN2A showed decrease in the hippocampi of both the animal models. In contrast to this data, the mRNA levels of *Grin2A* and *Grin2B* were high compared to controls indicating that there might be mechanisms such as regulation by miRNAs, by which the protein levels are being regulated (Figure 8A and B). Some previous studies have reported similar observations [53–55] in the hippocampus and/or prefrontal cortex. However there are also reports, which show no significant changes in the GluN2A/GluN2B expression in the hippocampus and/or prefrontal cor-tex [47,56]. This might be due to various factors such as differences in the age and sex of the animals used for the study, the treatment regime or the variation among the rat species. However, in the prefrontal cortex of the MK-801 treated animals, the protein expression of GluN2A and GluN2B was found to be high (data not shown) as reported by earlier studies in which changes were not found in the hippocampus [47]. The other schizophrenia related proteins that we studied, NRG1, BDNF and CaMKIIα, showed alterations that are divergent in the two models. In the MK-801 model, the transcript and protein levels increased whereas in the MAM model, the transcript levels were not significantly altered while the levels of the proteins were either unchanged or decreased indicating mechanisms involving miRNAs. The decrease in GluN2A and GluN2B proteins is consistent with NMDAR hypofunction in both the models while alterations in the levels of NRG1, BDNF and CaMKIIα proteins could cause dysfunction in synaptic regulation (Figure 7A and B).

To test whether translational arrest by miRNAs could be a possible mechanism for the downregulation of NMDAR subunit proteins, we estimated the levels of miR-148b, miR-223, miR-129-2, miR-296 and miR-137 that were found to target *Grin2A* and *Grin2B* in our *in vitro* experiments. It was found that one or both of the complimentary strands (5p/3p) of some of these miRNAs were significantly high in the hippocampus in both the doses of MK-801 treatment and in MAM treatment implying that these miRNAs might actually be regulating the NMDAR subunits. Interestingly, miR-223 that is known to lower the expression of GluN2B and protect the brain from neuronal cell death [27] and miR-129-2, remained unchanged, in the MAM model implying that they might not be regulating *Grin2B* in this condition.

Recent studies have focused on the variation of the 5p/3p ratio of miRNA strands, as in the case of miR-574, from tissue to tissue, during development and also during disease conditions depending on the regulatory roles to be performed [57,58]. We observed that 5p and 3p forms of miR-148b, miR-129-2, miR-296, miR-223 and miR-137 were co-upregulated in the MK-801 model. Intriguingly there was an inverse correlation of the expression levels of 5p and 3p strands of miR-148b and miR-137, in the MAM model. Such regulation of coexpression of the 5p/3p strands of miRNAs can either be through target mediated miRNA protection (TMMP) or though target RNA directed miRNA degradation (TDMD). TDMD has been observed in the neurons as a process for miRNA regulation [59–61]. It is possible that the changes in the expression levels of miR-148b, miR-129-2 and miR-296 are guided by the increased levels of *Grin2A* and *Grin2B* expression.

This is the first report, which shows that miR-296-3p, miR-129-2-3p and miR-148b-5p are involved in regulating the NMDA receptors. Previous studies have reported alterations in the miRNA levels in the MAM model by RNA sequencing [62,63] but our study involves extensive research on miRNAs and their role in regulating NMDAR in the MAM model.

In the MK-801 model, NMDAR is directly blocked by MK-801. It is known that MK-801 can inhibit NMDAR activity in the brain when administered systemically [64]. Sustained inhibition of NMDAR activity could elicit a feedback response of increase in expression of NMDAR subunit transcripts (Figure 10A) [65](Wilson et al., 1998). High transcript levels are translationally arrested by high levels of the NMDAR targeting miRNAs, leading to decreased translation resulting in lower protein expression. Reports have suggested that NRG1 can regulate NMDAR [66,67], but due to the binding of MK-801 to NMDAR, NRG1 may become functionally in-competent with respect to NMDAR regulation. We can also speculate that high levels of NRG1 transcript might be an effect of reduced NMDAR as these are closely associated in PSD region. In parallel, high levels of NRG1 may regulate BDNF, thus in-creasing its expression in the system [68]. We also found significant increase in CaMKIIα transcript levels in the 0.5 mg/Kg MK-801 treatment with not much change in protein levels, implying an active miRNA regulation. Bioinformatic analysis showed that miR-148b-5p could target CaMKIIα and hence we hypothesize that miR-148b-5p, which was significantly high in animals that received 0.5 mg/Kg MK-801 dose downregulated the protein levels of CaMKIIα. There is an increase in the CaMKIIα transcript levels with significant increase in the protein in the 1.0 mg/Kg dose of for MK-801 treated animals, which could be a direct/indirect effect of the hypofunction of NMDAR.

**Fig 10.**
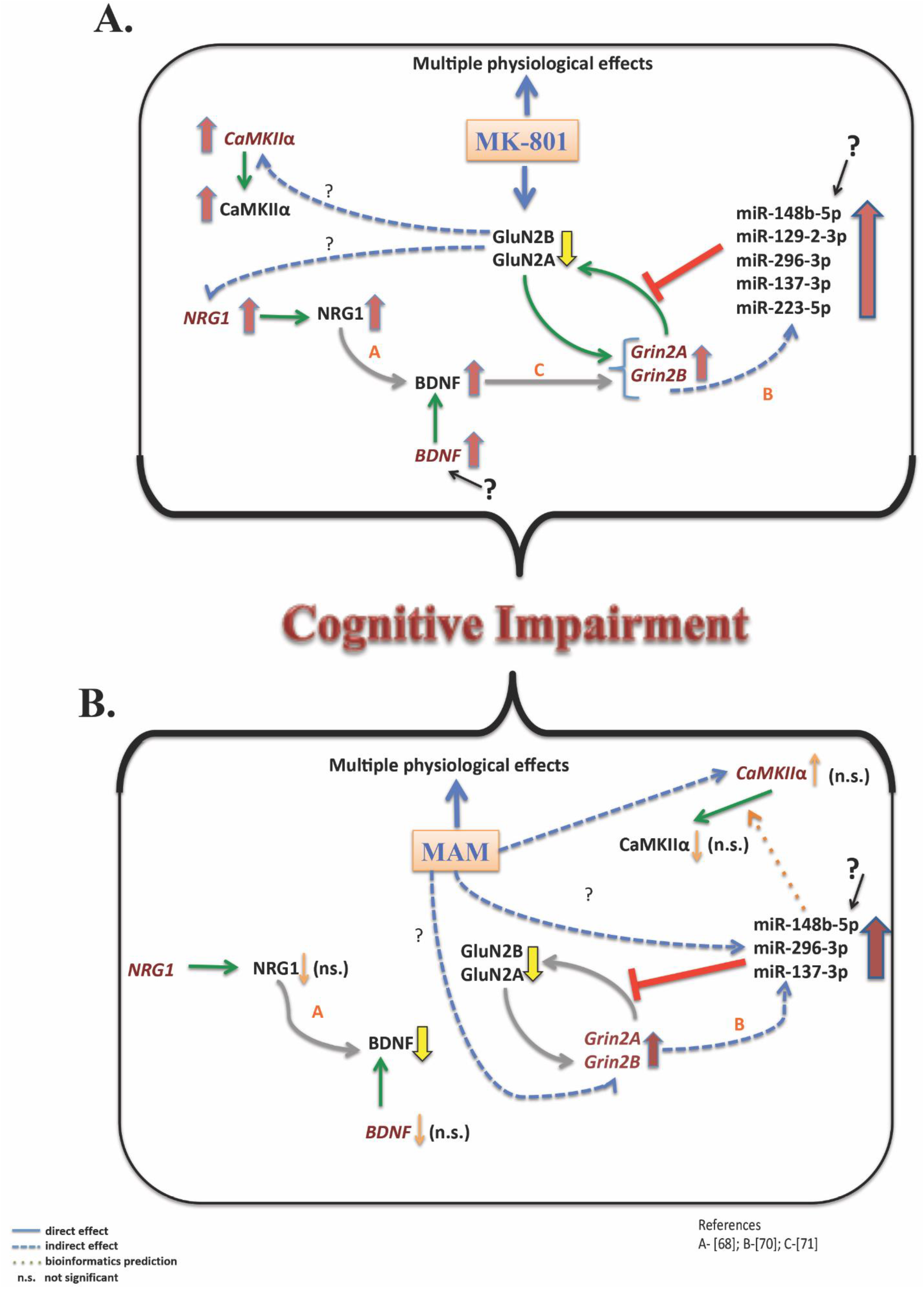
Schematic representation of the regulation of expression of NMDAR subunits-GluN2A and GluN2B and related proteins in MK-801 model and MAM model of schizophrenia. (A) In the MK-801 model the activity of NMDAR is low due to inhibition by MK-801 leading to increased *Grin2A* and *Grin2B* expression as a scaling response for replenishing the protein level. But due to translational arrest by the upregulated miRNAs such as miR-129-2-3p, miR-296-3p, miR-148b-5p and 3p, miR-223-5p and miR-137-3p, there is decrease in protein levels. The NRG1 and CaMKIIα levels are high, which might be a direct/indirect effect of the pharmacological action of MK-801. Increase in NRG1 levels could be leading to an increase in the BDNF level (Pandya and Pillai, 2014). (B) In the MAM developmental model, there is an abnormal neurodevelopment pattern with altered synaptic circuit leading to decrease in GluN2A, GluN2B and BDNF expression. There is an increase in the transcript levels of *Grin2A* and *Grin2B* probably due to a feedback mechanism but the levels of the miRNAs that target Grin2A and Grin2B are high leading to translational inhibition and downregulation of protein levels.

In case of the MAM model, the neurotoxin itself can affect neurogenesis in the CNS, which may result in altered expression of GluN2A and GluN2B (Figure 10B). The transcript levels of *Grin2A* and *Grin2B* are high either as a direct consequence of the action of MAM or as a response to reduced protein level, but the abundance of NMDAR targeting miRNAs arrest translation, leading to reduced protein production (Figure 10B). Bioinformatic analysis showed that miR-148b-5p could target CaMKIIα (data obtained using three or more prediction tools). We postulate that this miRNA might target and decrease the protein levels of CaMKIIα in the MAM model. From the q-PCR data, we compared the relative expression levels between both the models and found that miR-148b-5p is considerably higher (10 fold) in the MAM model than the MK-801 model and CaMKIIα mRNA concentration was more or less same in both the models. However the magnitude of increase of miR-148b-5p is significantly higher in the MAM model so that they could target CaMKIIα transcripts and successfully cause downregulation of the protein.

miR-137 is a well-studied miRNA in schizophrenia and it is known to regulate the GluN2A subunit of NMDAR. We observed an enormous increase in the miR-137-3p levels in the MAM model with respect to the control, indicating that miR-137-3p might be a major player in regulating *Grin2A* transcript and could also cause the schizophrenia phenotype in the MAM model. It will be interesting to check if other target proteins predicted for these miRNAs are indeed undergoing changes. Through bioinformatic analysis we also inferred that miR-296, miR-129-2 and miR-148b has other target proteins involved in synaptic plasticity such as the α-amino-3-hydroxy-5-methyl-4-isoxazolepropionic acid receptors (AMPAR) and voltage gated calcium channels (Suppl. Table 1).

Dysregulation of NMDARs have been associated with several disorders apart from schizophrenia, such as depression, Huntington’s disease and Alzheimer’s disease [69]. Pharmacological manipulation of NMDAR usually leads to side effects due to lack of specificity towards receptor subtypes among other reasons. Regulatory mechanism such as those involving miRNAs are targeted towards specific subunits and hence can achieve receptor subtype specificity thereby having excellent therapeutic potential. These miRNAs can either be used as biomarkers or as therapeutic agents for altering the levels of NMDAR and thus minimizing disease outcome.

## 4. Materials and Methods

### 4.1 Materials

Tris, ammoniumpersulphate (APS), phenylmethylsulfonyl fluoride (PMSF), 30% acrylamide, Tween-20, sodium dodecyl sulphate (SDS), β-mercaptoethanol, CaCl_2_, dithiothreitol (DTT), tetramethylethylenediamine (TEMED), protease inhibitor cocktail (PIC), dizocilpine (MK-801) and NaOH were from Sigma Chemicals, USA or from Amershampharmacia biotech, USA. Methylazoxymethanol acetate (MAM) was from FUJIFILM Wako Pure Chemical Corporation, Japan. LB agar, LB broth, ampicillin, agarose, Tween-20 and bovine serum albumin (BSA) were from GE Healthcare Life Sciences. Antibiotic-antimycotic, lipofectamine 2000 reagent, Dulbecco’s modified eagle medium (DMEM), Glutamax, trypsin, fetal bovine serum (FBS), Hank’s balanced salt solution (HBSS), Opti-MEM and Pierce BCA (bicinchoninic acid) protein assay kit were from Thermo Fisher Scientific. Restriction digestion enzymes and DNA ladders were from Fermentas. T4 DNA ligase enzyme was from New England Biolabs. mirVana kit from Invitrogen was used for the isolation of miRNAs. Mir-X™ miRNA First Strand Synthesis Kit was from Takara Co., Ltd., Dalian, China. High-Capacity cDNA Reverse Transcription Kit was from Applied Biosystems, USA. Prestained protein marker, Clarity ECL reagent, polyvinylidene difluoride (PVDF) and nitrocellulose membranes were from Bio-Rad, USA. Antibodies against GluN2B from Santacruz (Cat. No-sc9057) or Abcam (Cat. No-Ab65783), BDNF from Abcam (Cat. No-Ab108379), Neuregulin from Santacruz (Sc28916), CaMKII from CST (Cat.No 3357S) and β-actin from Sigma (Cat.No-A5316) were used.

### 4.2 Methods

#### 4.2.1 Bioinformatic analysis

The following miRNA prediction tools were used for mining the miRNAs that target the genes *Grin2A* and *Grin2B* - TargetScan (http://www.targetscan.org/vert_72/), miRanda (http://www.microrna.org/), Micro-Cosm (https://tools4mirs.org/software/mirna_databases/microcosm-targets/), Segal/PITA (https://genie.weizmann.ac.il/pubs/mir07/mir07_prediction.html) and miRDB (http://mirdb.org/). To alleviate false positives, prediction was accepted only if three or more tools gave consistent results. RNAhybrid tool (https://bibiserv.cebitec.uni-bielefeld.de/rnahybrid/) was used to check the stability of binding between the miRNAs and the 3’UTR of the genes *Grin2A* and *Grin2B* using minimum free energy as the criteria. The 3’UTR sequences of the genes were obtained using UCSC Genome browser (https://genome.ucsc.edu/) and the sequences of the miRNAs were obtained using miRbase.

#### 4.2.2 DNA constructs

To generate the constructs of the 3’UTR and the pre-miRNAs, primers were designed for the sequences obtained from UCSC database. The 3’UTR of Grin2A (NM_012573.3) and Grin2B (NM_012574.1) were amplified from rat genomic DNA and were cloned into Xho1 and Not1 sites of the dual luciferase vector, psiCHECK2 (Promega) where the 3’UTR of Renilla luciferase is replaced with the 3’UTR of Grin2A or Grin2B. The firefly luciferase cDNA present in the plasmid was used for normalization. Similarly, pre-miRNAs were also amplified from rat genomic DNA and were cloned into the intronic region of the DsRed gene in the expression vector, pRIPM. DsRed served as a marker for checking the transfection efficiency. The clones were confirmed by restriction digestion and sequencing. The empty pRIPM vector was used as a control for transfection experiments. The vectors, psiCHECK2 and pRIPM were obtained as gifts from Dr. Jackson James and Dr. Ani V Das, Rajiv Gandhi Centre for Biotechnology, India and Dr. Beena Pillai, Institute of Genomics and Integrative Biology (IGIB), India respectively. The primers used for cloning the pre-miRNA constructs and the 3’UTRs are listed in Table 2.

**Table 2.**
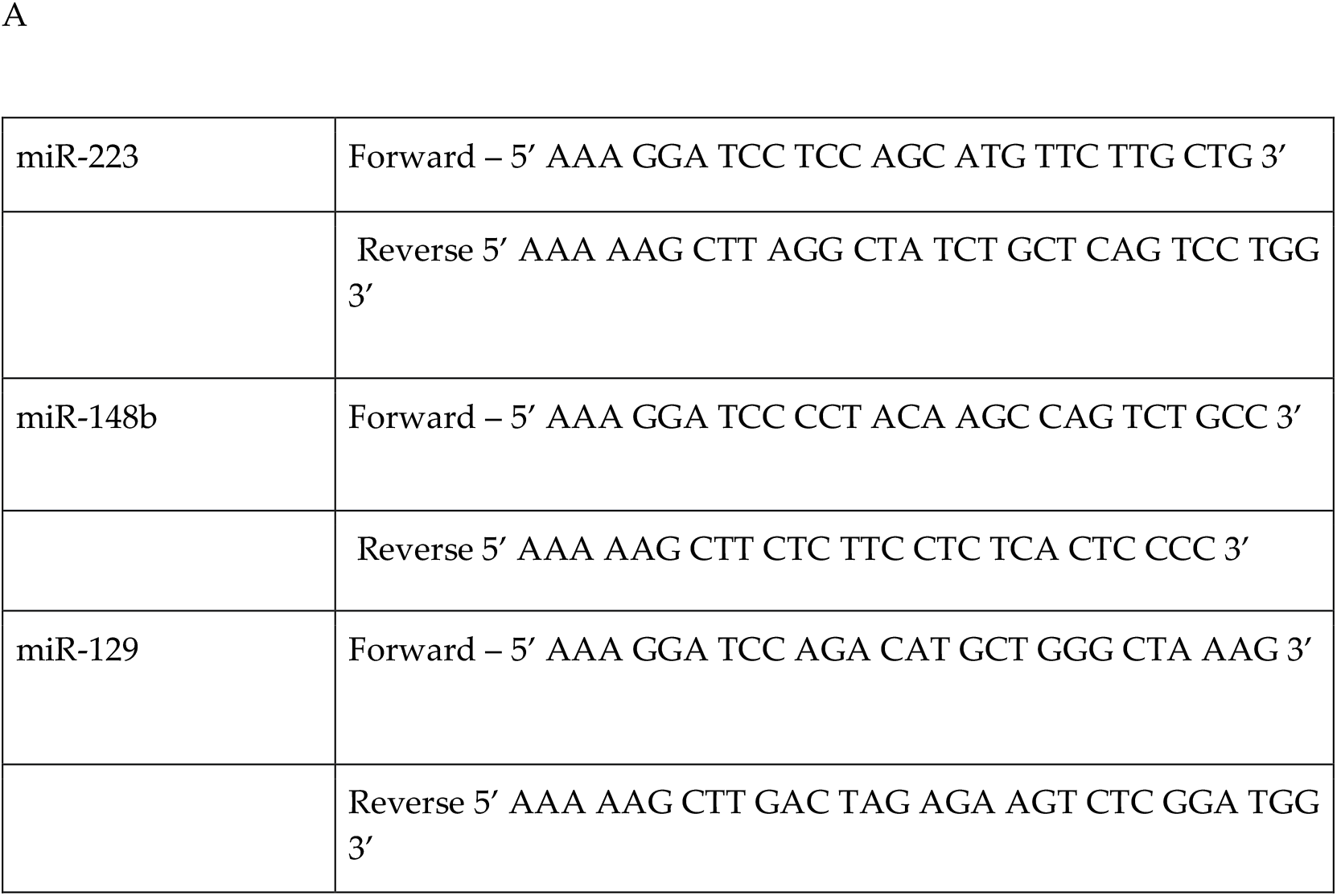
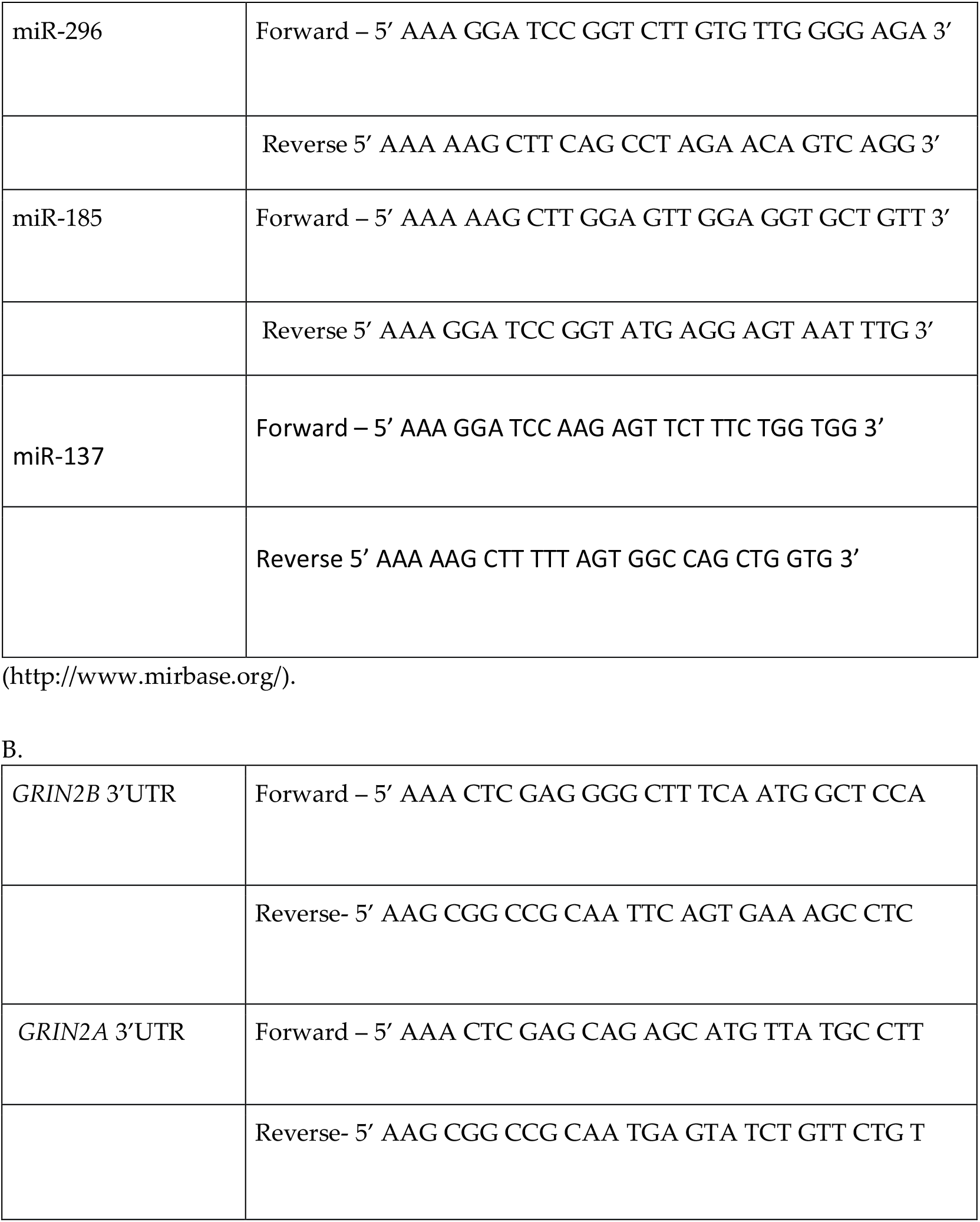
Primers used for cloning pre-miRNAs and 3’UTRs. (A) Primers used for cloning the pre-miRNAs into the pRIPM vector were designed using the UCSC genome database. (B) Primers used for cloning the 3 ‘UTR for *Grin2A/2B* genes into the Promega psiCHECK2 vector were designed using the UCSC genome database

#### 4.2.3 Luciferase assay

Human Embryonic kidney-293 (HEK-293) cells were maintained in DMEM containing 10% FBS with 100 units/mL of penicillin and 100 μg/mL of streptomycin at 37°C in a humidified environment of 95% O2 and 5% CO2.

About 1 × 105 cells were seeded in each well of a 24 well plate. The cells were cotransfected with psiCHECK2 vector containing the *Grin2A* or *Grin2B* 3’UTR and pRIPM vector carrying the pre-miRNA in 1:1 ratio using Lipofectamine in OptiMEM medium. The empty pRIPM vector along with 3’UTR vector was used as control. This mix was incubated at room temperature for 30 min and was added to the cells. Complete media change was given after 6-8 hrs of incubation. It was observed that the transfection efficiency was around 85-90% by 24-48 hrs following transfection (Suppl. Figure 1). The cells were harvested after 48 hrs and luciferase assay was performed using the dual reporter kit (Promega) according to manufacturer’s instructions.

#### 4.2.4 Animals

Animals were housed separately with standard chow, water ad libitum and 12 hr light-dark cycle. Wistar rat embryos of embryonic day 18-19 (E18-19) were used for primary neuronal culture preparation. Adolescent male Wistar rats (40-45 day old, 100-150g) were used for creating the MK-801 model. For the MAM model, pregnant female Wistar rats were injected with MAM and the offsprings, both male and female, were used as the schizophrenia model. All the procedures were approved by the Insti-tutional Animal Ethics Committee (IAEC) of Rajiv Gandhi Centre for Biotechnology and followed the rules and regulations prescribed by the Committee for the Purpose of Control and Supervision of Experiments on Animals (CPCSEA), Government of In-dia.

##### 4.2.4.1 Primary neuronal cultures and transfection

Primary cultures of hippocampal neurons were prepared from E18-E19 rat embryos from pregnant dams [20,21]. The hippocampi were dissected from the brain, digested with trypsin and the dissociated cells after sufficient washes were seeded on-to plates, which were precoated with poly-D-lysine (100 μg/ml) and laminin (1μg/ml). The neurons were grown in neurobasal medium supplemented with 1% B27 supplement, 2 mM glutamax and 100 μg/ml penicillin-streptomycin at 37°C in a hu-midified environment of 95% O2 and 5% CO2. The medium was partially changed every two days. The cultures were maintained up to days in vitro (DIV 14).

Temporal profile of expression of Grin2B transcript was studied using cultures from DIV 3 to DIV 14. The seeded cells were scrapped on the required days (DIV 3, 5, 7, 9, 12 and 14) using 1X PBS, centrifuged, snap frozen and kept at −80°C. The cell pellets were subjected to RNA isolation followed by qPCR analysis.

Neuronal transfections were conducted using Lipofectamine 2000 reagent on DIV 7. For each well of a 24-well plate, a total of 1-1.5 μg of pre-miRNA plasmid was used with 1:50 dilution of lipofectamine 2000 in neurobasal medium. The empty vector pRIPM was used as the control for the experiments. The transfection mix was incubated at room temperature for 20-30 min and was further diluted two-fold in neu-robasal medium. Neurons were incubated with this mix for 4-6 hrs and then the medi-um was completely replaced with neurobasal medium. The cells were fixed with 4% paraformaldehyde on DIV 10.

###### 4.2.4.1.1 Immunocytochemistry

Hippocampal neurons that were transfected with pRIPM vectors containing the pre-miRNAs on DIV 7 were fixed with 4% paraformaldehyde in PBS on DIV 10. After washing, the neurons were incubated with primary antibody-rabbit anti-GluN2B (1:500, Abcam) overnight at 4°C followed by incubation with fluorophore conjugated secondary antibodies (Alexa Fluor 488). Z-stack images were acquired using a confocal microscope (Nikon Eclipse Ti A1R with NIS Elements V.4.0 soft-ware). Maximum projection images are presented and were used for ROI based image quantitation.

##### 4.2.4.2 Treatment regimes for the animal models

###### 4.2.4.2.1 MK-801 model using male Wistar rats

MK-801 (0.5 mg/Kg and 1.0 mg/ Kg) (n=6 per group) was injected intraperitoneally, once a day for 5 days between 2 pm and 5 pm every day to male Wistar rats of postnatal day 40-45. The control group (n=6) received the same volume of saline. This was followed by a five-day drug washout period and subsequent behavioral analysis such as open field test (OFT), novel object recognition test (NORT), object location test (OLT) and Morris water maze (MWM) test. The rats were sacrificed by cervical dislocation and hippocampi were dissected [22]. These were used for bio-chemical analysis such as western blotting and quantitative real time PCR.

###### 4.2.4.2.2 MAM model using male and female Wistar rats

MAM (20 mg/Kg) was administered intraperitoneally to pregnant female Wistar rats on gestational day 17 (GD 17)/ embryonic day 17 (E17). The control dams received saline injection on E17. The pups were weaned on postnatal day 30 and were placed in single separate cages. Both male and female pups were used for this model. The behavioral tests were initiated on/after P45-P50 (n=10-13 for MAM group, n=10 for control group). The experimental rats were sacrificed by cervical dislocation after the behavioral tests and the dissected hippocampi were used for biochemical analyses.

##### 4.2.4.3 Behavioural analysis

###### 4.2.4.3.1 Open field test (OFT)

The animals were transported to the behavior chamber at least one hour prior to the start of each experiment to acclimatize the animals to the test facility environment. The time spent in the central and peripheral zone and the total distance travelled by the animal in an open-field box (70 cm × 70 cm × 40 cm, lbh respectively) in 10-11 min were measured by video recording using either a Panasonic CCTV camera (WV CP 500) or LifeCam studio webcam by Microsoft (Q2F-00013) placed above the arena. The box was cleaned using 70% alcohol between the trials. All analyses of the video files were carried out using Noldus EthoVision XT software (Noldus Infor-mation Technology, Wageningen, The Netherlands).

###### 4.2.4.3.2 Novel object recognition test (NORT) and object location test (OLT)

NORT was conducted in an open field arena (70 cm × 70 cm × 40 cm lbh respectively) with 2-3 different kinds of non-toxic objects. The objects used were generally consistent in height, volume and color but are different in shape and appearance. During habituation, the animals were allowed to explore an empty arena. Recording during habituation is used as data for OFT. Twenty-four hours after habituation, each animal was subjected to training by exposing them for 5 min to the familiar arena in which two identical objects were placed. At 1 hr and 24 hrs after training, the rats were subjected to test by allowing them to explore the open field in the presence of one of the familiar object and a novel object for 5 min. The box and the objects were thoroughly cleaned using 70% alcohol between the trials. MK-801 treated animals were subjected to test at 1 hr after training and MAM treated animals were subjected to test at 1 hr and 24 hrs after training. The time spent by the animal exploring each object was scored using the video file. The discrimination index (DI) and recognition index (RI) were calculated as follows.

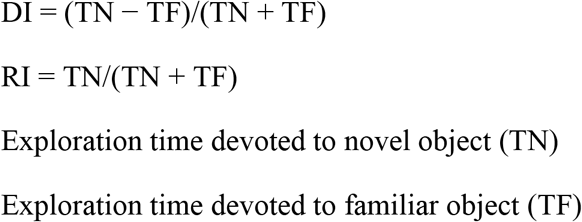

In OLT, the animal was allowed to explore the familiar arena with two identical objects for 5 min. For the test session, after one hour, one of the objects was retained at the same familiar location as the training session (non-displaced object), while the other was moved to a new location (displaced object). The rat was placed back into the arena for 5 min and exploration of the objects was measured. DI and RI were calculated.

###### 4.2.4.3.3 Morris water maze (MWM) test

The water maze test was performed using an opaque white tank, 170 cm in diameter and 80 cm in height with a circular platform (around 20 cm diameter) in one of the quadrants of the maze. The maze was filled with water till the platform was submerged and the water was made translucent by adding milk. Rats were placed in water and were trained to find the hidden platform. The training consisted of 5 trials of 60 seconds each with intertrial intervals of 20-30 seconds per day for 5 consecutive days. Recordings were done by the video camera placed above the arena and analysis was done using Noldus EthoVision XT software. The time the animal takes to reach the platform (escape latency) was noted and the average escape latency for the 5 trials on each day was calculated for each animal.

The animals were sacrificed after the MWM test and the brains were immediately removed, dissected, snap frozen and stored in −80°C.

##### 4.2.4.4 Analysis of hippocampal tissues by quantitative real time polymerase chain reaction (qPCR)

The frozen hippocampal tissues were made to a fine powder by homogenizing after freezing in liquid nitrogen. The miRNA and total RNA were extracted by mir-Vana miRNA isolation kit. The cDNA of the miRNA was synthesized using the Mir-X^™^ miRNA First Strand Synthesis Kit. The cDNA for the genes was generated using High-Capacity cDNA Reverse Transcription Kit. Quantitative real-time PCR (qPCR) was done for different genes and miRNAs using the SYBR green system and relative expression levels in each sample were measured using the 2^ΔCT value. For genes, actin was used as internal reference and for miRNAs, U6-sno was used as internal reference. The primers used for the genes were KiCqStart SYBR green primers (Sigma) and the primers for miRNAs are listed in Table-2.

**Table-3.**
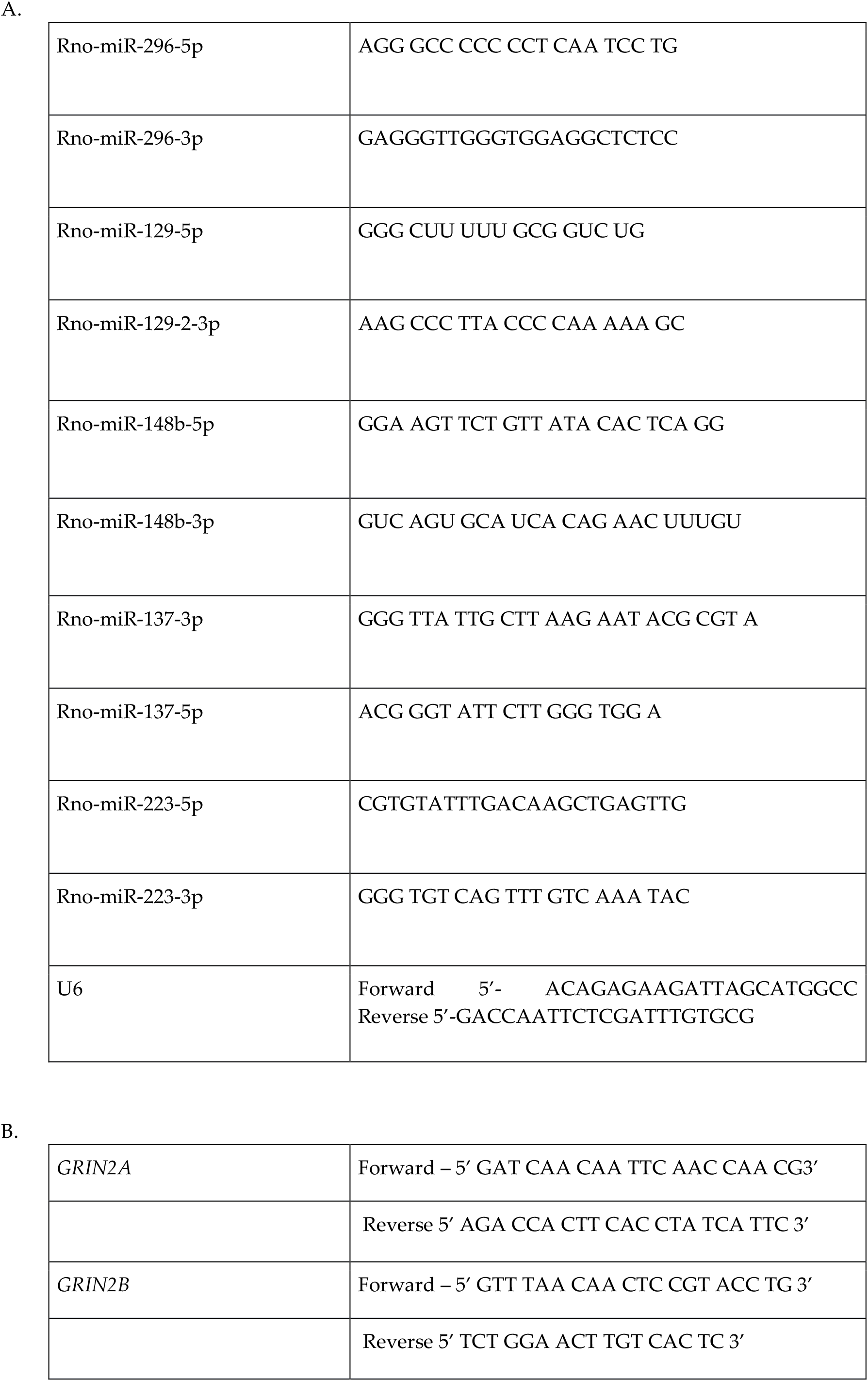
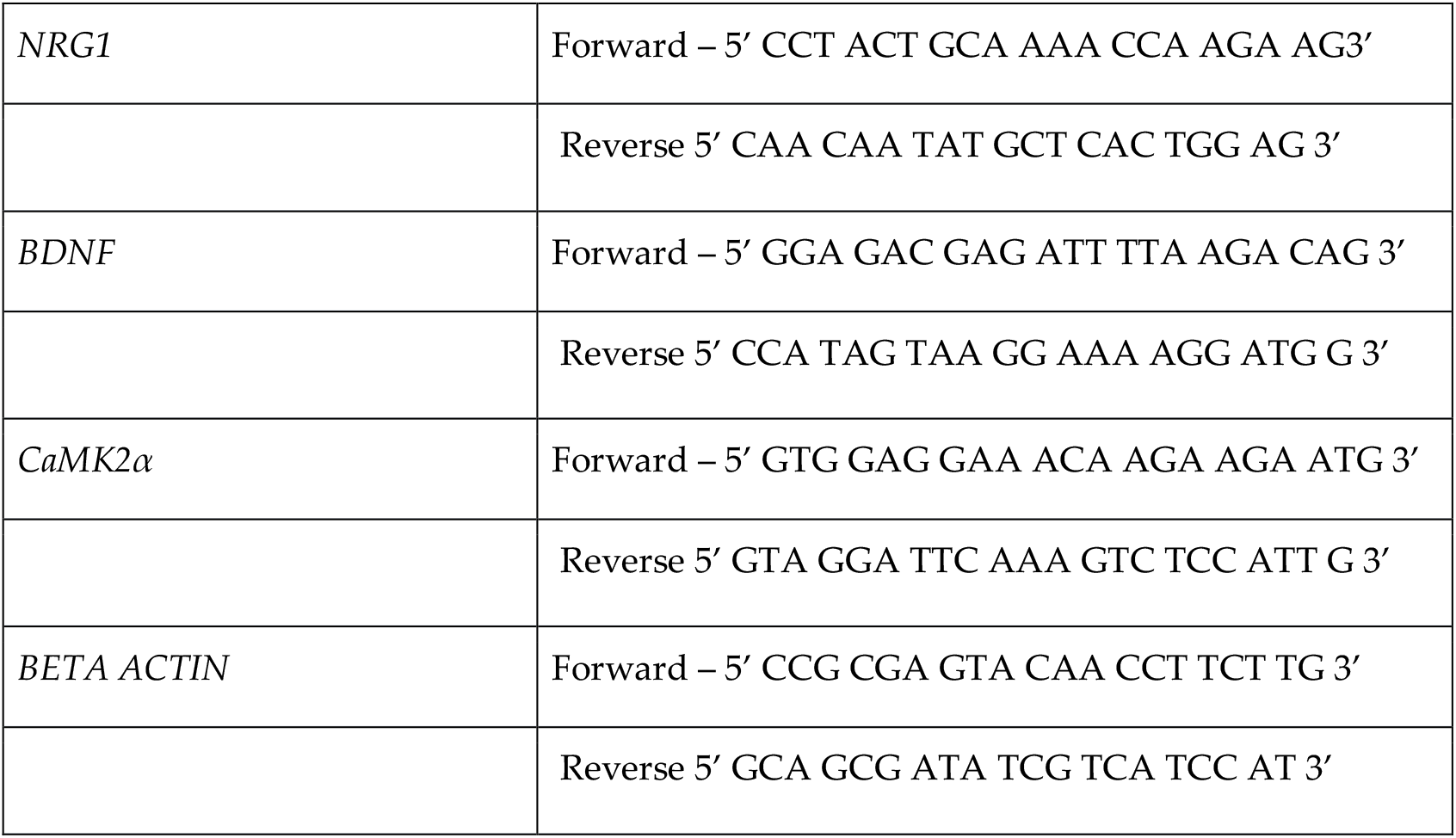
Primers used for real time PCR of miRNAs and genes. (A) Primers used for amplifying the transcript of the miRNAs. (B) Primers used for amplifying the transcript of the genes.

##### 4.2.4.5 Western blot analyses of hippocampal tissues

The tissues were finely ground and were solubilized in lysis buffer [50 mM Tris-HCl, pH 7.4, 150 mM NaCl, 1% SDS, 1% NP-40, 5 mM EDTA, 50 mM β-glycerophosphate, 5 mM sodium orthovanadate, 0.2 mM phenylmethylsulphonylfluoride (PMSF), 1 mM dithiothreitol (DTT), 10 mM staurosporine and 1X complete protease inhibitor cocktail]. The protein concentrations were estimated by BCA method. Around 60-100 μg protein of each sample was subjected to SDS-PAGE and was transferred to polyvinylidene difluoride (PVDF) or nitrocellulose membrane. The membranes were blocked with 5% bovine serum albumin (BSA) and then probed with the following primary antibodies - mouse anti-GluN2A (1:1000, Millipore), rabbit anti-GluN2B (1:1000, Abcam), mouse anti-beta actin (1:3000, Sigma), rabbit anti-neuregulin (1:1000, Santa cruz), rabbit anti-BDNF (1:1000 Abcam), and mouse anti-CaMKIIα (1:1000, Cell Signaling Technologies). Incubation with primary antibody was carried out overnight. After washing, blots were incubated with horseradish peroxidase (HRP)-conjugated secondary antibodies (1:3000 or 1:10000, Sigma) and were then developed in Chemidoc gel apparatus (Biorad, USA) using Clarity ECL reagent. The intensity of each band was quantitated using ImageJ software. The data were normalized to band intensity of β-actin.

#### 4.2.5 Statistical Analysis

The data is reported as mean ± standard deviation (SD) and is based on a minimum of three independent experiments/replicates. Luciferase assay data (n=3-4) was analyzed using unpaired parametric Student’s t test to compare between each miRNA group and the corresponding control group. For immunocytochemical analysis [n (Total no. of fields quantified in 3-4 experiments)=15-22], the average fluorescence intensities for the untransfected and transfected cells for all the replicates were calculated and were analyzed using unpaired Student’s t test. For MK-801 animal model, data from the behavioural experiments (n=6 per group) and the biochemical measurements (n=4-6 per group) such as western blotting and qPCR, were statistically analyzed using the one-way ANOVA followed by Dunnett’s post hoc test for multiple comparison between the control and the two treatment groups (MK-801 −0.5 mg/Kg and 1.0 mg/Kg) except for MWM experiments. Data analysis for MAM model was done using the unpaired parametric Student’s t test. Data from the MWM tests for MK-801 model and MAM model were analyzed using repeated measures two-way ANOVA as there are two variables - the time and the groups. This was followed by Dunnett’s post hoc test for the MK-801 model and Sidak’s post hoc test for the MAM model for multiple comparison between the control group and the treatment group/groups. The analysis for males and females were initially done for the behaviour experiments of the MAM model. But no major difference was found between males (n=6 for saline, n=5 for MAM)) and females (n=4 for saline, n=8 for MAM)) and hence the data has been combined and represented together for both sexes in behaviour experiments (n=10-13 per group). This study did not address the sex differences in the biochemical measurements (n=3-4 per group) such as qPCR and western blotting for the MAM model. The statistical analysis for the behaviour experiments for both the models were completely blinded. The data was analyzed using the software, GraphPad Prism version 8 (8.4.2). Statistical significance was set at p <0.05.

## 5. Conclusions

In summary, we show that miR-129-2, miR-148b and miR-296 interact with *Grin2A* and *Grin2B* subunits of the NMDAR by using bioinformatics, luciferase assay and overexpression in primary hippocampal neurons. Our results from *in vivo* studies in rats treated with MK-801 or MAM, show that GluN2A and GluN2B proteins are downregulated with concomitant upregulation of their transcript levels indicating transcriptional and translational regulation, consistent with the NMDAR hypofunction model of this disorder. We unveiled that miR-148b-5p, miR-137-3p and miR-296-3p have significantly high expression in both the animal models indicating these miRNAs might be critical regulators of NMDAR. miR-129-2-3p and miR-223-5p showed upregulation only in the MK-801 model. There was an inverse expression pattern of the 5p and 3p forms of miR-148b and miR-137 in the MAM model. This study has revealed novel and significant roles for certain miRNAs in regulating the expression of NMDAR, which also appear to participate in the pathophysiology of schizophrenia. Further investigation of the mechanism of these miRNAs might pro-vide promising avenues in therapeutics.

## Associated content

Additional tables and figures are given in the supporting information.

## Acknowledgments

We are grateful to Dr. Jackson James and Dr. Ani V Das for providing the luciferase plasmid psiCHECK2 (Promega) and pRIPM plasmid. We thank Mr. K.C. Sivakumar for technical advice in bioinformatics. We thank Dr. Mayadevi, Dr. Mantosh Kumar, Dr. Lakshmi K and Ms. Amata Boban for their support and advice.

