## Supplementary Information for "Novel microRNAs targeting NMDA receptor subunits in animal models of schizophrenia"

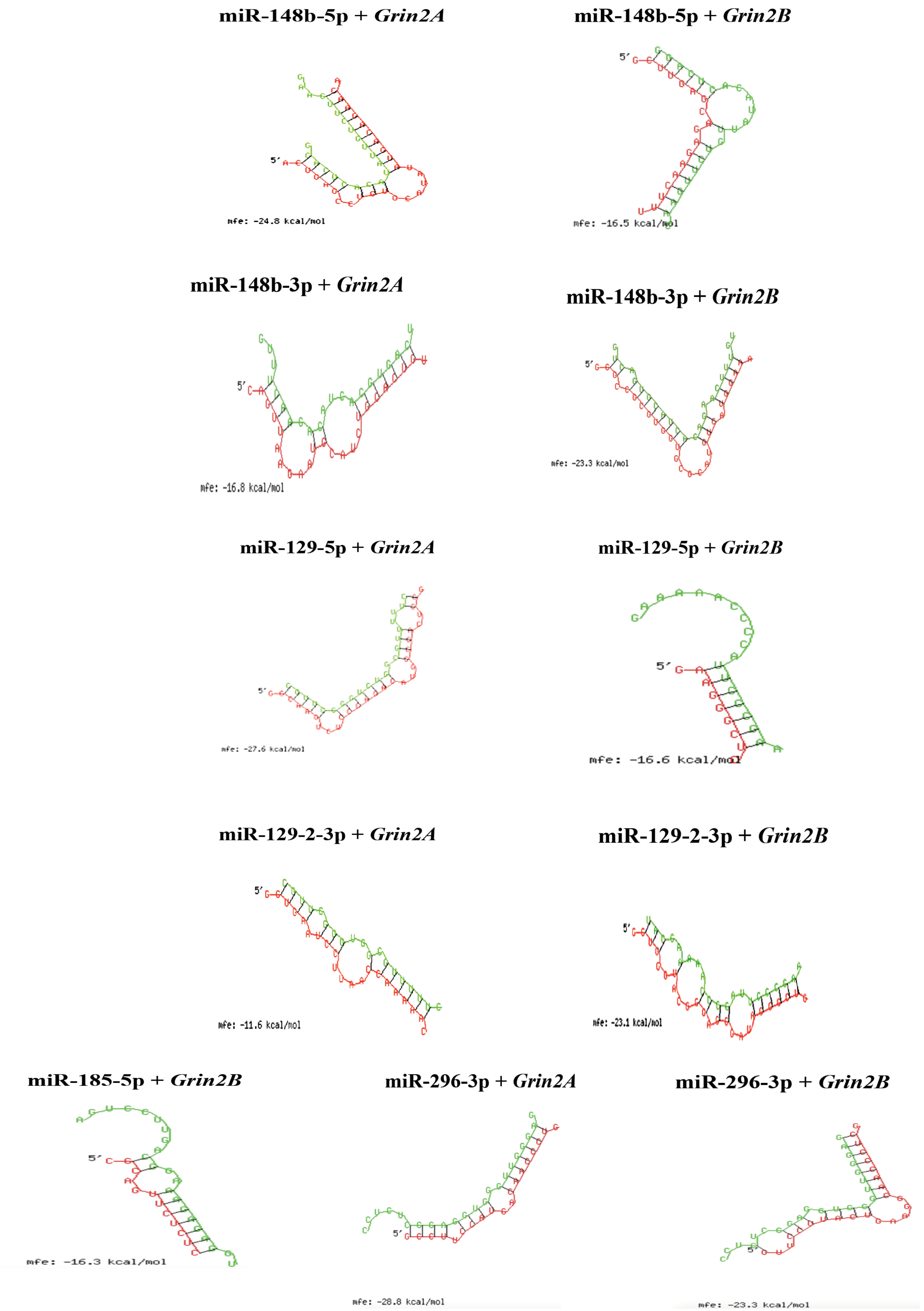


Suppl. Figure 1. RNA hybrid tool with predicted hybridization of the miRNAs (green sequence) with *Grin2A* 3'UTR and *Grin2B* 3′ UTR (red sequence). Minimum free energy (mfe) required for hybridization is indicated for each of the miRNA-mRNA binding complexes

A

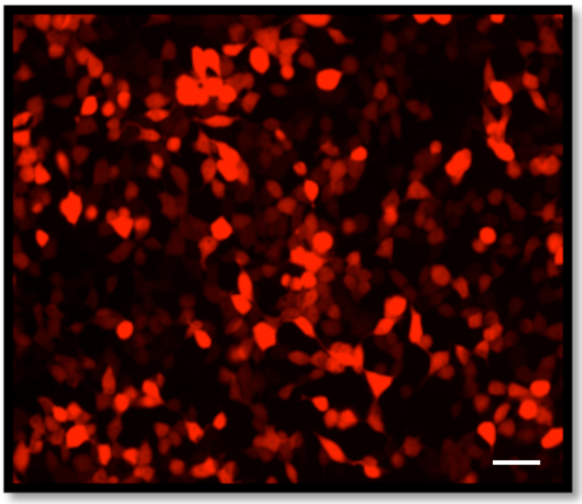

B

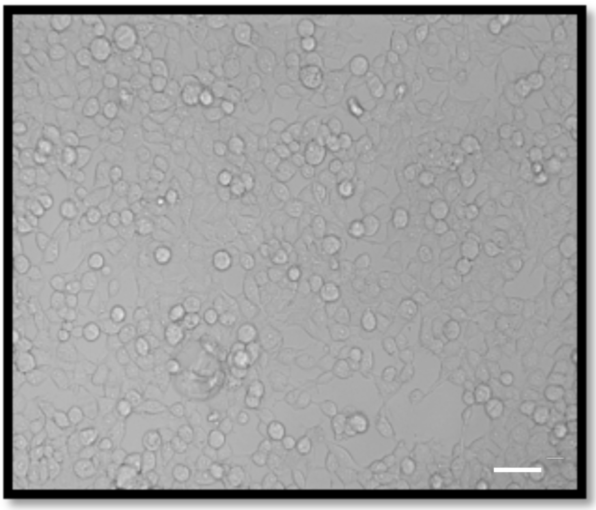

Suppl. Figure 2 Representative image of transfection of the pRIPM vector with miR-148 in HEK-293 cells. Image was captured after 48hrs of transfection. (A) Fluorescence image of the DsRed marked transfected cells. (B) Brightfield image of the cells. Scale bar: 20µm

| | miR-129-2 | miR-148b | miR-296 |
|---|---|---|---|
| *CACNA1C* | ✓ | ✓ | ✓ |
| *CACNA1I* | ✓ | ✓ | ✓ |
| *CACNB2* | ✓ | ✓ | ✓ |
| *ZNF804A* | ✓ | ✓ | ✗ |
| *ErbB4* | ✗ | ✗ | ✓ |
| *GRIA1* | ✗ | ✗ | ✓ |
| *GRIN2A* | ✓ | ✓ | ✓ |
| *GRIN2B* | ✓ | ✓ | ✓ |
| *GRIN3A* | ✓ | ✓ | ✓ |
| *GRM1* | ✓ | ✓ | ✓ |
| *GRM3* | ✓ | ✗ | ✓ |
| *NRG1* | ✗ | ✗ | ✓ |
| *NRG3* | ✗ | ✓ | ✗ |
| *CaMK2α* | ✓ | ✓ | ✗ |

Suppl. Table 1. Selected miRNAs can target candidate genes of schizophrenia. Bioinformatic analysis showed that these miRNAs could also target the genes involved in the pathology of schizophrenia. CACNA1C (Calcium Voltage-Gated Channel Subunit Alpha1 C); CACNA1I (Calcium Voltage-Gated Channel Subunit Alpha1 I); CACNB2 (Calcium Voltage-Gated Channel Auxiliary Subunit Beta 2); ZNF804A (Zinc Finger Protein 804A); ERBB4 (Erb-B2 Receptor Tyrosine Kinase 4); GRIA1 (Glutamate Ionotropic Receptor AMPA Type Subunit 1); GRIN3A (Glutamate Ionotropic Receptor NMDA Type Subunit 3A); GRM1 (Glutamate Metabotropic Receptor 1); GRM3 (Glutamate Metabotropic Receptor 3); NRG3 (Neuregulin 3)
